# Database for extended ligand-target analyses (DELTA): a new balanced resource for AI applications in drug discovery

**DOI:** 10.1101/2025.05.22.655476

**Authors:** Arianna Pisati, Agnese Pozzi, Federico Giuntini, Alessandro Pedretti, Giulio Vistoli

## Abstract

We here present the DELTA resource, a database including balanced and annotated datasets of ligands for about 500 therapeutically relevant targets specifically collected for developing AI-based predictive models. For each target, DELTA comprises an optimized protein structure plus 200 experimentally tested ligands equally distributed between active and inactive molecules. All ligands are prepared by considering unspecified isomeric elements and combining semi-empirical calculations with MD simulations to explore their conformational space. The so-collected molecules allowed extended analyses of both ligands and targets, and the study presents some preliminary results. The performed analyses revealed that on average active ligands are larger than inactive molecules, while possessing a similar polarity. The scaffold analysis emphasized the expected and crucial role of aromatic systems, even though with some relevant differences between active and inactive molecules. Moreover, similar targets often show conserved binding sites and there is a limited but not negligible relationship between the similarity of binding sites and ligands suggesting that similar pockets tend to bind rather similar ligands. Finally, the collected biological data also allowed the analysis of the polypharmacological profile of the ligands endowed with more than one biological value. Most ligands bind two or three targets with diverse activities and almost always the bound targets belong to the same biological class. All the collected data are available for download at delta.unimi.it.

## Introduction

Over the past few years, the drug discovery process benefitted from the huge increase of available biological and chemical data as well as from the endless development of ever more performant predictive approaches. The combination of these two factors markedly enhances the role of *in silico* methods (especially of AI-based approaches) in accelerating drug design and discovery [1,2]. Nevertheless, the increased relevance of AI algorithms should be paralleled by the development of tailored data resources to successfully exploit them, while most available resources were designed and collected for different applications [3].

Indeed, the available resources can be roughly subdivided into two classes. The first group comprises annotated and curated datasets of high-quality structures of ligand-target complexes to be primarily used to support and validate the development of both docking engines and scoring functions (e.g. see refs. [4,5]). All these resources usually include an equal number of ligands and protein targets, and the collected complexes are almost exclusively experimentally resolved structures. The second group comprises datasets in which few active ligands for a given target are dispersed within a huge number of inactive compounds. They are primarily collected to support the development and assessment of innovative strategies in virtual screening campaigns (e.g. see refs. [6,7]). Usually, these resources combine experimentally determined active molecules and in silico generated inactive compounds (the so-called decoys).

PDBbind [8,9] represents the prototype of the first class of resources: the last 2024 release collects about 33,600 high-quality complexes annotated with affinity values to empower both structural and energetic predictions [10]. Also, BioLip2 [11] is a curated database of high-quality biologically relevant ligand-protein complexes, which combines the structures from PDB with other information taken from literature or other online resources. BioLip2 is constantly updated, and the release of January 2025 includes about 93,500 entries. Lastly, the recently reported PLINDER resource represents the largest and most annotated database of curated protein-ligand interactions extracted from about 163,000 resolved structures [12].

The resources for the second group can be exemplified by MUV [13], DUD [14] and DUD-E [15] tools, which collect datasets of ligands for 18, 40 and 103 targets, respectively. Each dataset includes a variable number of experimentally determined active ligands (e.g. the number of actives is, on average, equal to 650 in DUD-E) plus a huge number of decoys to reach a percentage of actives equal to about 3%. Decoys are molecules generated in silico by matching the physicochemical properties of the actives while preserving a satisfactory structural diversity [16]. The recent LIT-PCBA resource includes 21 targets for which both active and inactive compounds are experimentally determined, and various resolved structures are available for each target. In this way, LIT-PCBA represents an improved tool for unbiased assessments of both ligand and structure-based virtual screening campaigns [17].

All mentioned resources were extensively exploited and are finding many applications in rational drug design, even though they do not possess the optimal features to be applied for comprehensive classification analyses using AI-based approaches. To do this, an optimal data resource should include, for each target, only experimentally tested compounds equally subdivided between active and inactive molecules. The included compounds should not compulsory be numerous, but the selected ones should suitably cover the chemical space of the ligands for the considered target. In addition, such a resource should comprise balanced datasets for many therapeutically relevant targets to enable extended comparative analyses. Ideally, there should be experimentally resolved structures for all considered targets to allow reliable structure-based studies.

Based on the above, we here present DELTA (Database for Extended Ligand-Target Analyses) a novel resource which contains balanced datasets for about 500 therapeutically relevant human targets as derived from TTD database [18]. For each target, DELTA includes a dataset of 200 experimentally tested ligands equally subdivided between active and inactive compounds. Even though larger balanced datasets could have been collected for many targets, we decided to limit the selected ligands to 200 for all targets to have homogeneous and equinumerous datasets. These harmonized datasets are enough to develop reliable classification predictors and allow comprehensive benchmarking analyses. For all targets, DELTA includes reliable 3D structures, which are almost exclusively experimentally resolved, to enable reliable structure-based simulations. The structures of both ligands and targets are refined and prepared to be easily exploited in various calculations. The DELTA resource is available at delta.unimi.it.

## Methods

### Selection of the targets

The therapeutically relevant target proteins were collected by considering the list of the targets for both marketed drugs (623 proteins) and drugs in clinical trials (1037 proteins) as compiled by the 2022 update of the Therapeutic Target Database (TTD) [18,19].

The structures of the targets were collected by applying the following criteria:

1. If more than one resolved structure was available in the Protein Data Bank (PDB), the structure was selected by considering: (a) the presence of a bound ligand within the active site, (b) the resolution of the structure and (c) the lack of unresolved gaps.
2. In the case of homomultimeric proteins, only a single monomer was selected according to its structural quality and completeness.
3. If no resolved structure was available, homology models as generated by using SWISS-MODEL [20] were accepted only when the similarity with a resolved structure was greater than 0.8.
4. If no homology structure was available, ab initio models as generated by deep learning-based AlphaFold algorithm [21] were accepted only when the average pLDDT score of the model was greater than 0.7.

The so-selected protein structures were retained only if a suitable number of active and inactive ligands were available (see below). In this way, 477 protein structures were collected which represent about one third of all the targets included in the two considered lists from TTD. The selected structures are equally distributed between targets for marketed drugs and in clinical trials.

### Preparation of the protein structures

The resolved structures were prepared by removing crystallization chemicals and non-relevant co-crystallized molecules using the VEGA ZZ suite of programs [22]. Unresolved gaps and loops were filled by modelling them, if required, by homology techniques using the ChimeraX’s interface of MODELLER [23, 24]. As mentioned above, the homology models were generated by using the SWISS-MODEL resource and the template was selected by considering both the sequence similarity and the coverage. The *ab initio* models were downloaded from the AlphaFold Protein Structure Database [21] by applying the above-mentioned filter based on the overall confidence of the retrieved structures. All the experimental and theoretical protein structures were carefully verified by checking and repairing (when necessary) the chirality of residues, the geometry of peptide bonds and possible steric clashes between side chains. The hydrogen atoms were added by using VEGA ZZ and, to be compatible with physiological pH =7.4, Glu, Asp, Lys and Arg residues were considered ionized, while His and Cys were maintained neutral by default. A specific script was developed to automatically check the correct assignment of the hydrogen atoms to each residue. The so-obtained protein structures were minimized using NAMD [25] by keeping the backbone atoms fixed to preserve the overall (predicted or resolved) folding.

The binding sites of the selected protein structures were analysed by using the MatGAT application [26]. The binding sites were defined as the residues with at least an atom within a 10 Å radius sphere around the binding site center. If the original PDB structure of the protein contains a co-crystallized ligand, the binding site center was calculated as the center of the pocket formed by residues with at least an atom within 8 Å of the ligand. In the other cases, literature was examined to identify the residues of the binding site and if no result was found, the key residues were predicted using COACH [27]. Once the residues were identified, the point at the center of the selected residues was considered as the center of the binding site.

The structural similarity of the proteins was evaluated by the TMscore which is computed by the software TMalign as a metric for estimating topological similarity of proteins [28]. Scores below 0.17 correspond to structurally unrelated proteins, while scores higher than 0.5 generally correspond to proteins with similar folding. For all protein pairs, the highest TMscore (the TMscore relative to the shortest protein of the pair) was used to populate the similarity matrix. The distance matrix was then computed by difference from the similarity matrix (Distance = 1 – TMscore) and was fed to K-medoids PAM clustering [29] using the the kmedoids python library [30].

### Selection of the ligands

For each target, ligands with experimentally determined activity (measured in terms of IC50, XC50, EC50, AC50, Ki, Kd, or Potency) has been retrieved by ChEMBL [31] and PubChem [32] along with an extensive literature search. All the biological activities are expressed as pChEMBL values. The collected molecules have been filtered excluding inorganic or metalloorganic compounds as well as molecules with a molecular weight exceeding 1000 Da. To differentiate between active and inactive ligands, a threshold has been established corresponding to a pChEMBL value of 5.5. Ligands with a pChEMBL > 5.5 were categorized as actives, whereas those with a value lower than or equal to 5.5 were deemed inactives. When a balanced dataset of 200 molecules could not be collected due to strongly imbalanced source data, the threshold for the target was dynamically defined as the difference between the pChEMBL average and its standard deviation.

A KNIME workflow has been developed to automatize the preparation and the standardization of the ligands’ datasets. ln detail, the workflow (a) separates active and inactive ligands according to the defined threshold; (b) retrieves the SMILES string for each compound from the relative resource; (c) clusters the ligands and selects 100 active and 100 inactive molecules as diverse as possible by using the diversity picker as implemented in RDKit [33].

### Preparation of the ligand structures

After defining the datasets and collecting the corresponding SMILES strings, the first step in ligand preparation involved the generation of stereoisomers. The defined stereocenters were preserved, while all possible stereoisomers were generated for the unspecified stereocenters. For the sake of simplicity, molecules with more than 4 unspecified stereocenters were limited to 16 generated stereoisomers. The E/Z geometric isomerism was treated by using the same strategy, while the tautomeric forms were disregarded. With the collected SMILES strings for the ligands, potential duplicates were eliminated to ensure the generation of consistent structures for each dataset. The generation of the geometric isomers and stereoisomers and the check for duplicates were carried out by using the RDKit library [33]. In this way, 104757 structures have been overall collected.

The 3D structures have been generated and ionized according to physiological pH = 7.4. The entire database has been carefully checked, and erroneously generated structures have been manually corrected. All the so-prepared structures underwent a single point PM7 semi-empirical calculation by MOPAC (keywords = “PM7”, “1SCF”, “Precise”, “GEO-OK”, and “SUPER”) [34], which allows calculating accurate atomic charges plus a set of stereo-electronic parameters. Furthermore, these semi-empirical calculations serve as an additional check for the structural accuracy since MOPAC is unable to terminate its calculations when involving wrong structures.

The structure of each ligand was minimized by NAMD and the resulting structures underwent a heating process from 0K to 500K for 100 ps. Then, a molecular dynamics simulation was performed at 500K for 500 ps. Finally, each frame of the MD trajectory underwent minimization using NAMD, and, after frames clustering, the best structure of the five (when available) clusters with the lowest energies were selected to collect the final structures of the ligands.

For the reported analyses, the collected ligands were profiled by calculating four sets of descriptors and fingerprints: (1) 26 physicochemical descriptors as computed by VEGA ZZ; (2) the Morgan fingerprints (1024 digits) [35]; (3) 19 stereo-electronic descriptors as calculated by PM7-based MOPAC calculations and (4) 119 various descriptors as provided by the RDKit tool [33]. Before performing PCA or similarity analyses, all descriptors were normalized by Standard Scaler (as implemented in scikit-learn). Likewise, all reported PCA and t-SNE analyses [36] were performed by using scikit-learn [37].

## Results

### DELTA in numbers

As reported in Methods section, DELTA comprises balanced datasets for 477 therapeutically relevant targets. For each target, DELTA includes the 3D structure of the protein plus 200 experimentally tested active (100) and inactive (100) ligands. Due to the promiscuity of many collected ligands (see below), DELTA overall includes 78938 unique ligands annotated with 95400 biological data. The database arrives to 104757 different structures when considering unspecified geometric and stereo-isomerism. In detail, a little less than half of the ligands (35882) include at least one chiral centre and about half of these chiral molecules include only one stereocenter. Chiral compounds are almost equally distributed between active and inactive molecules. Geometric isomerism is markedly less frequent and only 16664 ligands can exist in more than one geometric isomer. The rather limited number of added stereoisomers (25819) indicates that most isomeric elements were specified in the retrieved ligand structures. Among the 477 collected target structures, a vast majority is represented by experimentally resolved structures (426) plus 37 homology and 14 ab initio models.

### Analysis of the ligands

A first analysis was focused on the possible overall differences between active and inactive ligands. For this purpose, the 477 datasets of ligands were split into 954 sets by separating active and inactive molecules. For each set, the average values for 26 physicochemical and geometrical descriptors, as computed by VEGA ZZ, underwent a PCA analysis. Figure 1 shows the resulting 2D plot, in which the two principal components (PC1 and PC2) are displayed with convex hulls. While considering the expected merging of active (red points) and inactive (blue points) datasets, one may note that these points are not homogeneously distributed along the X axis, since active red points are clearly more abundant for positive PC1 values and, vice versa, inactive blue points are concentrated in the negative region of PC1. In contrast, PC2 seems not to have a discriminant effect since active and inactive sets are rather evenly distributed along the Y axis. As schematized in the inset of Figure 1, the active sets with PC1 > 0 almost double those with PC1 < 0 (280 vs. 197) and the opposite is observed for the inactive sets with PC1 < 0 (320 vs. 157). When analysing the features mostly contributing to the PC1 values (see Table A1), all descriptors with the highest loadings are related to molecular size (e.g., molecular volume, surface, weight and related descriptors), while PC2 is mostly determined by descriptors related to the polarity (see Table A2, e.g. PSA, dipole moment and number of H-bonding groups). Thus, Figure 1 suggests that the active compounds are, on average, larger than the inactive ones while having a comparable polarity.

**Figure 1.**
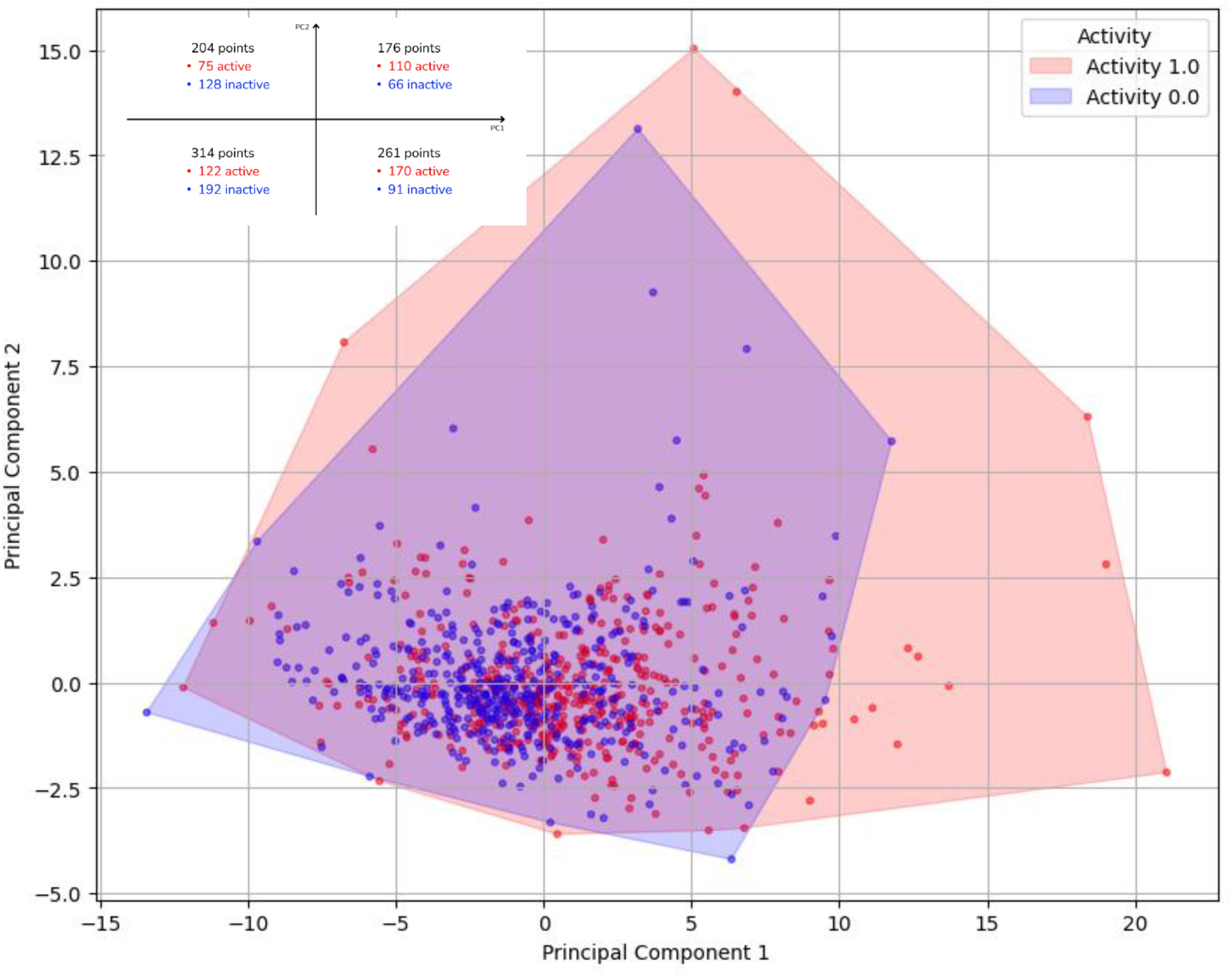
PCA analysis of 26 VEGA ZZ-based physicochemical descriptors for the 954 datasets as obtained by separating the active and inactive molecules of each target. The first two displayed principal components explain 71% of the overall variance. The inset schematizes the distribution of active and inactive sets in the four quadrants based on the PC1 and PC2 values.

To better investigate the differences between active and inactive datasets, the PCA analysis was repeated by considering and combining different sets of descriptors, but no other set of features provided the same explained variance reached by the already described VEGA ZZ-based descriptors (see Table A3).

While being less clear, a similar behaviour was observed when applying the PCA analysis to the VEGA ZZ-based descriptors on all ligands taken singularly (molecules active on one target, but inactive on other one, have been filtered out for this PCA analysis). The 2D plot of Figure 2 shows the two corresponding principal components (PC1 and PC2) with convex hulls and reveals a greater abundance of active ligands (red points) with PC1 > 0 and of inactive compounds (blue points) with PC1 < 0 (as detailed in the inset). Also here, PC1 is mostly determined by descriptors related to molecular size thus corroborating the above discussed considerations (see Tables A4 and A5 for loadings on PC1 and PC2, respectively).

**Figure 2.**
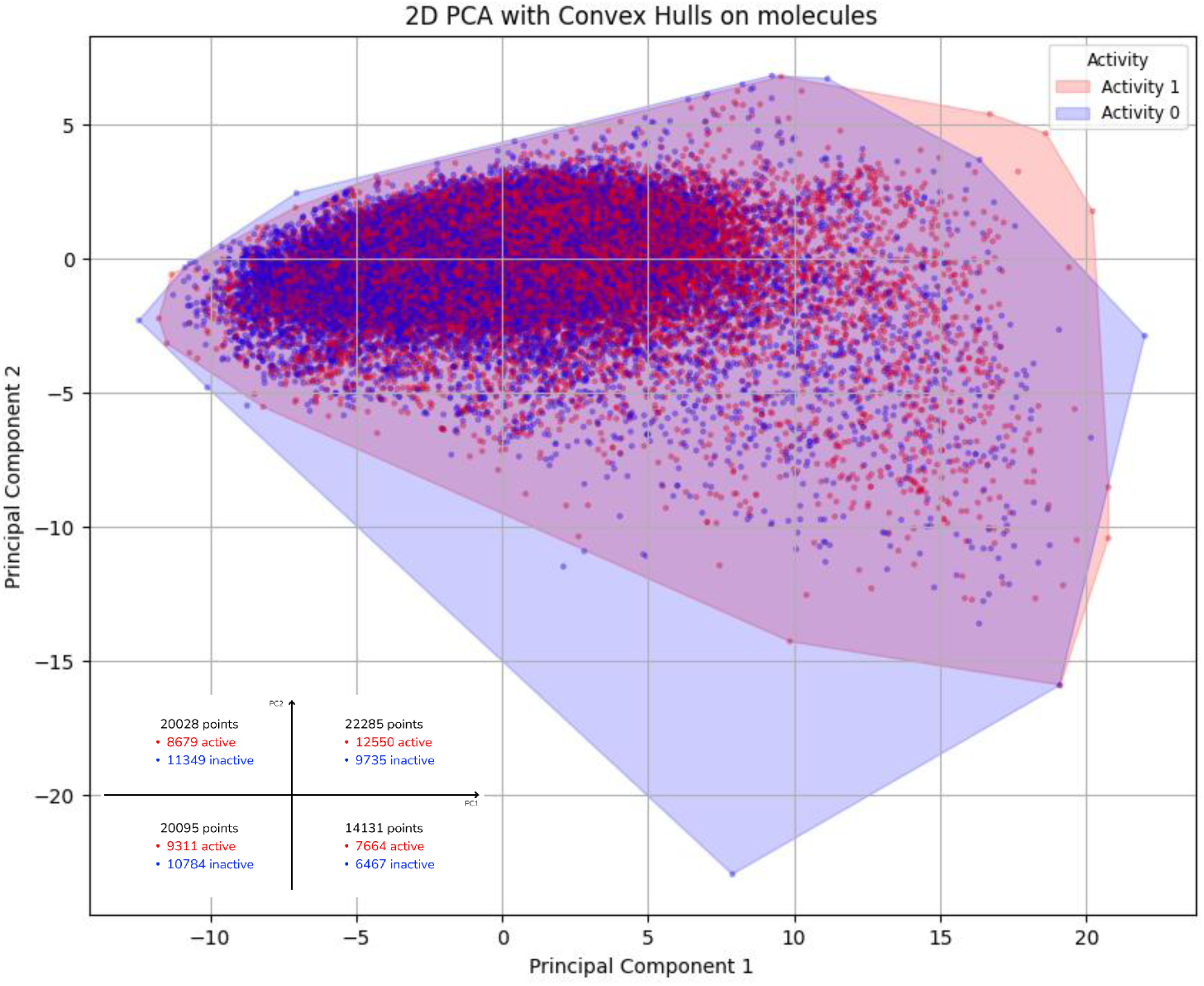
PCA analysis of 26 VEGA ZZ-based physicochemical descriptors for the individual ligands. The inset schematizes the distribution of active and inactive ligands in the four quadrants based on the PC1 and PC2 values.

Concerning the chemical space covered by the collected ligands, the analysis of Murcko scaffolds [38] revealed a certain degree of diversity between active and inactive compounds with the former being structurally more heterogenous. In detail, the active molecules include a higher number of scaffolds (22942 vs. 19517) and only 5109 scaffolds are found in both active and inactive ligands (molecules, that are active on one target, but inactive on other one, have been filtered out). Table 1 shows the 20 most frequent scaffolds (based on count) for active and inactive ligands.

**Table 1.**
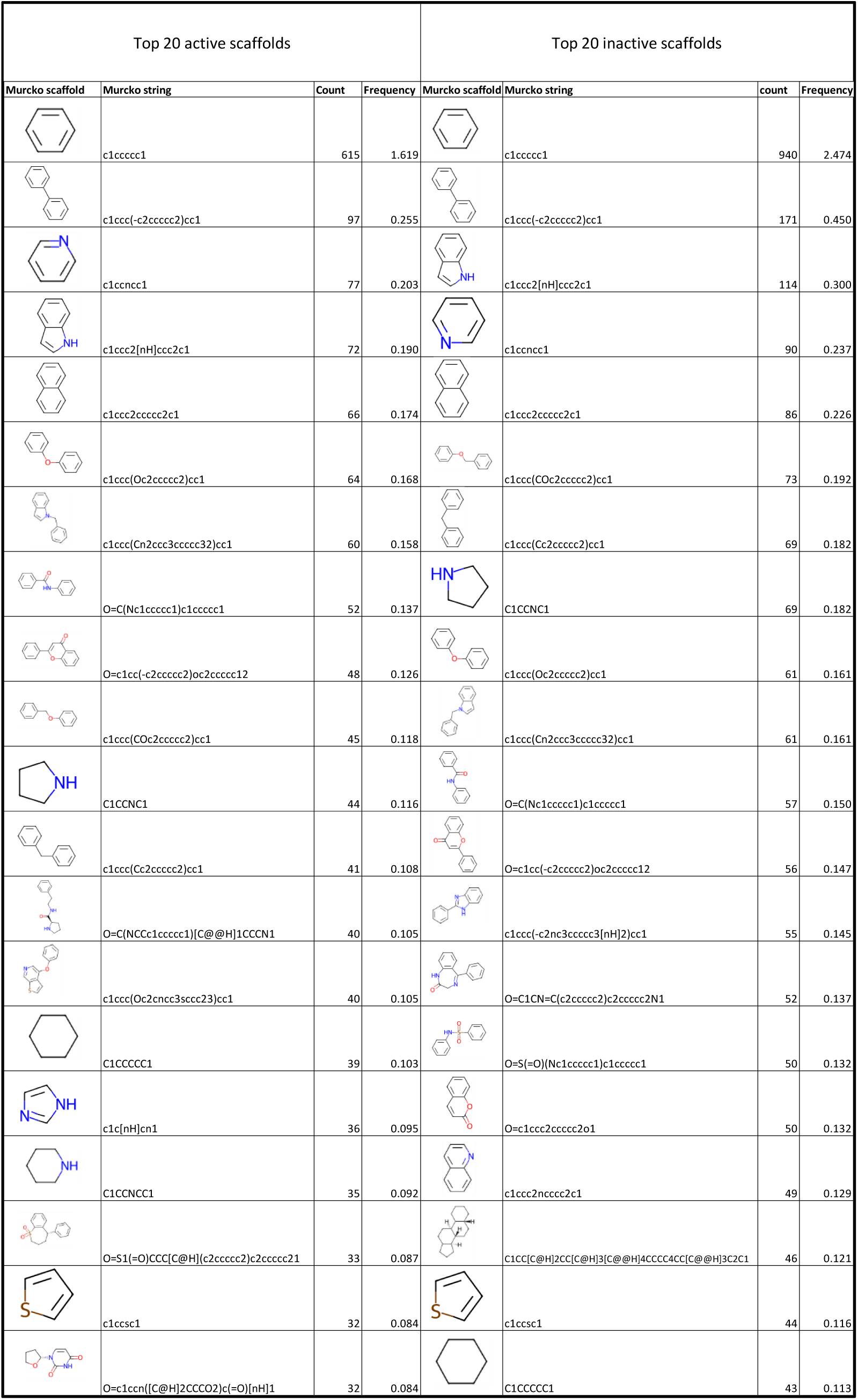
The 20 most frequent scaffolds (based on count) for active (on the left) and inactive (on the right) ligands. The frequency is reported as percentage values.

Apart from the expected abundance of aromatic systems, which similarly occupy the top of both rankings, marked differences are seen for the N-including rings. For example, piperazine, imidazole and pyrimidinedione are present only in the ranking of active compounds, while quinoline and benzodiazepine systems only among the top inactive scaffolds. Also, pyrrolidine is more frequent in the inactive compounds, while pyridine and indole are similarly abundant in both sets.

Next, Table A6 compiles the scaffolds which are present at least 10 times only in the active or only in the inactive compounds. While avoiding a systematic analysis of the found scaffolds, Table A6 reveals that the top active scaffolds include at most two aromatic moieties with only two exceptions, while 9 top inactive scaffolds out of 17 (including the top five moieties) include more than two aromatic groups. Also, the hydrolysable groups appear to be more frequent in the inactive scaffolds (9 *vs*. 3) and this difference might also find a pharmacokinetic explanation. Taken together, the scaffold analysis confirms the expected major role of the aromatic systems in all collected molecules and emphasizes that active ligands must possess a proper balance between lipophilicity and solubility and thus molecules including too many aromatic rings might not be active.

## Analysis of the targets

Figure 3 shows the distribution of the collected targets based on their biological class and reveals that most targets are composed by enzymes followed by receptors. According to EC classification, the two most represented classes of enzymes include the transferases and the hydrolases (with 128 and 94 proteins, respectively). Regulators, transporters, ion channels and nuclear hormone receptors are represented by a limited number of cases, the most populated class being the last one with 16 targets. Figure A1 reports the same pie charts as subdivided into targets of marketed drugs or in clinical trials and highlights that the relevance of receptors is more marked in the targets for marketed drugs compared to those for drugs in clinical trials, and vice versa for the enzymatic targets which are markedly more abundant in drugs in clinical trials.

**Figure 3.**
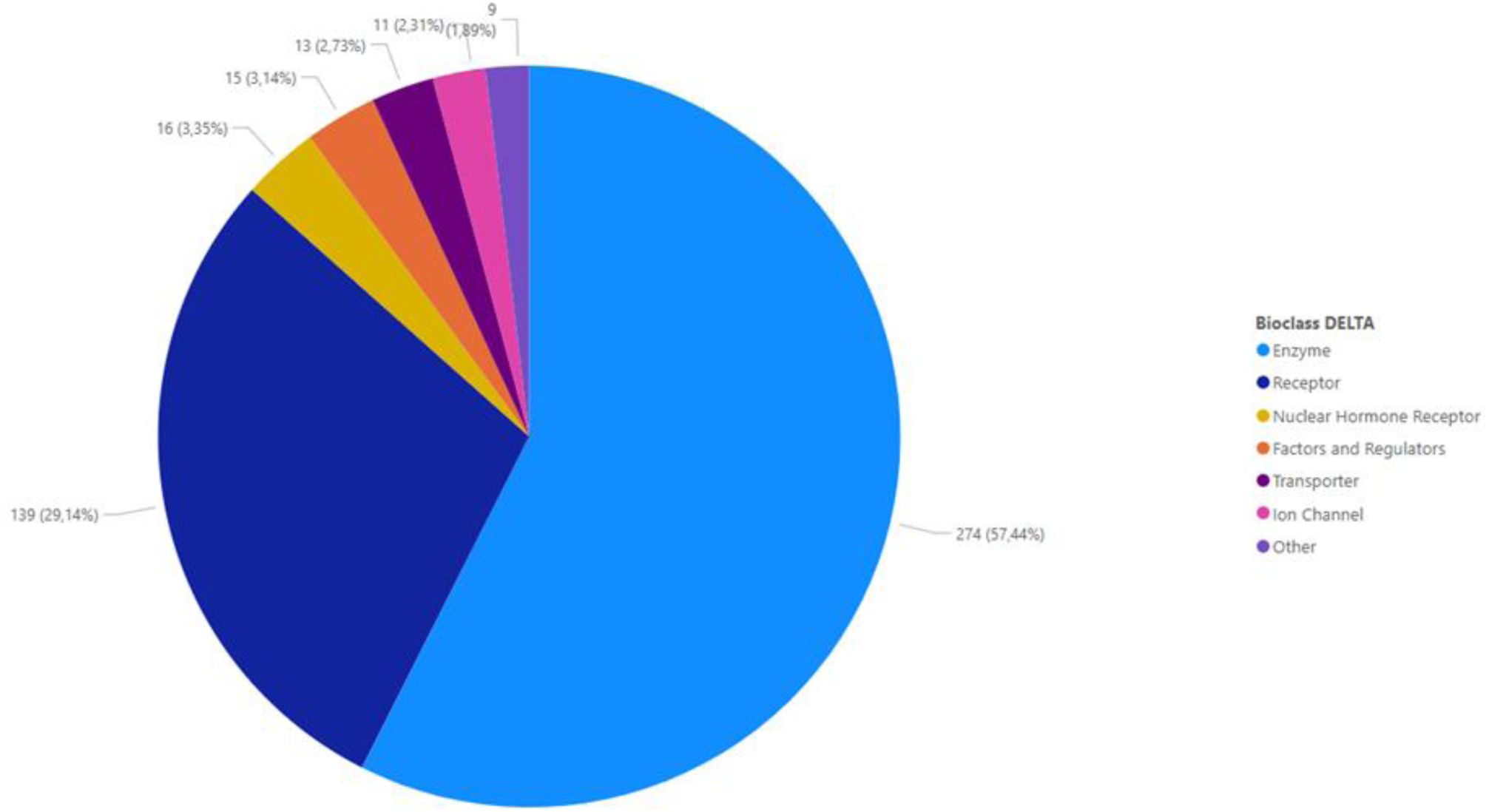
Classification of all 477 selected targets based on their biological class.

Figure 4 classifies the selected targets according to their therapeutic applications based on the International Classification of Diseases (11^th^ Revision) [39]. Notably, the only two classes with more than 100 targets are involved in anticancer activity or in CNS diseases. Well represented therapeutic applications (more than 60 targets) are also those involved in infective, metabolic, digestive, dermatologic and cardiocirculatory diseases. When comparing the classifications for marketed drugs or in clinical trials (data not shown), one may observe a markedly greater relevance of CNS targets in marketed drugs and of anticancer targets for drugs in clinical trials. Such a difference parallels what was observed for the biological classes since receptors are mostly involved in CNS diseases, while enzymes represent common anticancer targets.

**Figure 4.**
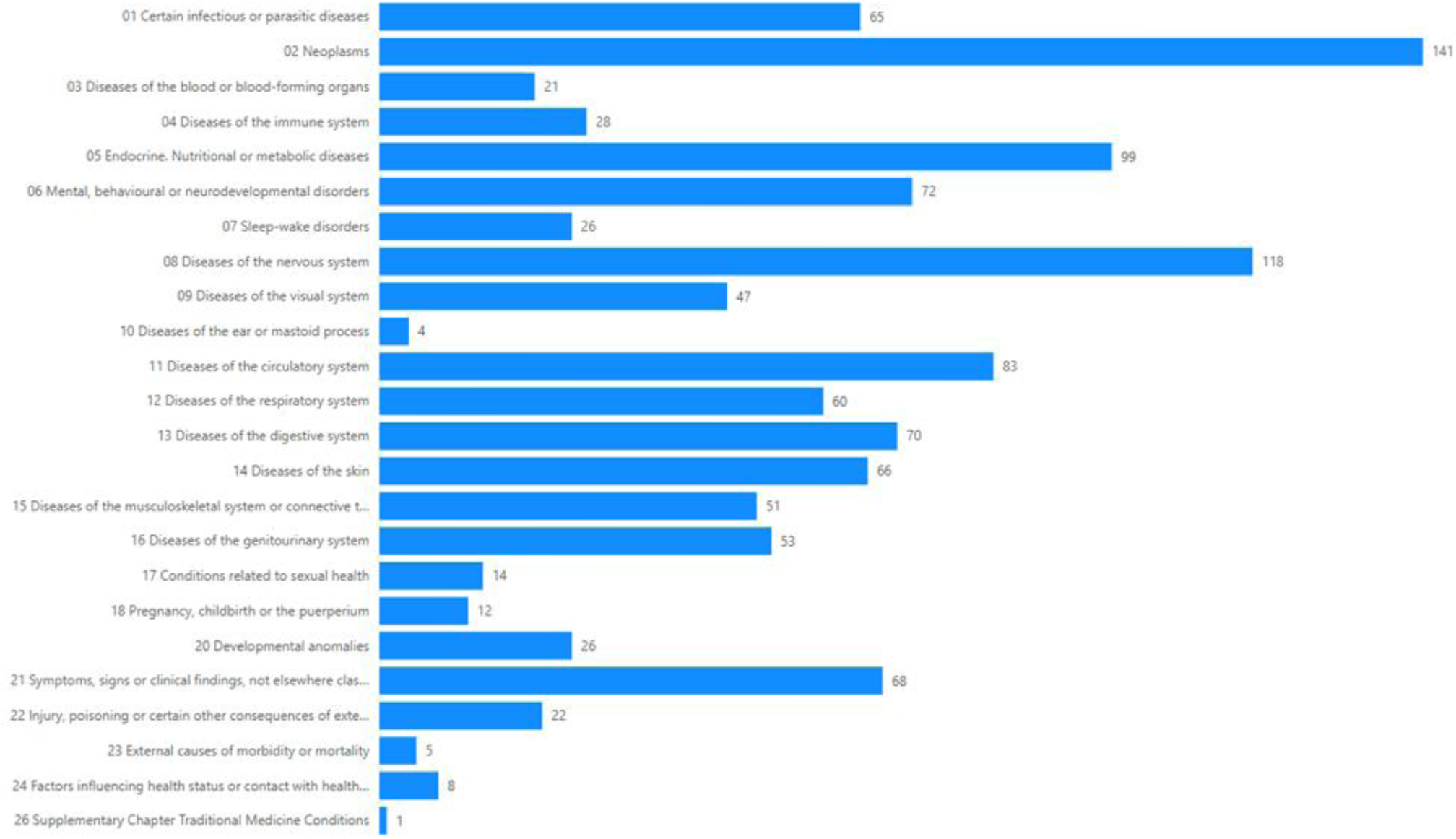
Classification of all 477 selected targets based on ICD-11 chapter.

Next, attention was focused on the relations between binding sites and protein structures. To simplify this analysis, the proteins were clustered based on their TMscore by using the K-medoids PAM clustering algorithm that identifies 20 clusters (as described under Methods). The MatGAT sequence similarity, as calculated for the binding sites only (see Methods), was utilized to populate the corresponding distance matrix (Distance = 100 – Similarity) which underwent t-SNE analysis [36]. Figure 5 displays the two t-SNE dimensions as coloured per K-Medoids clusters.

**Figure 5.**
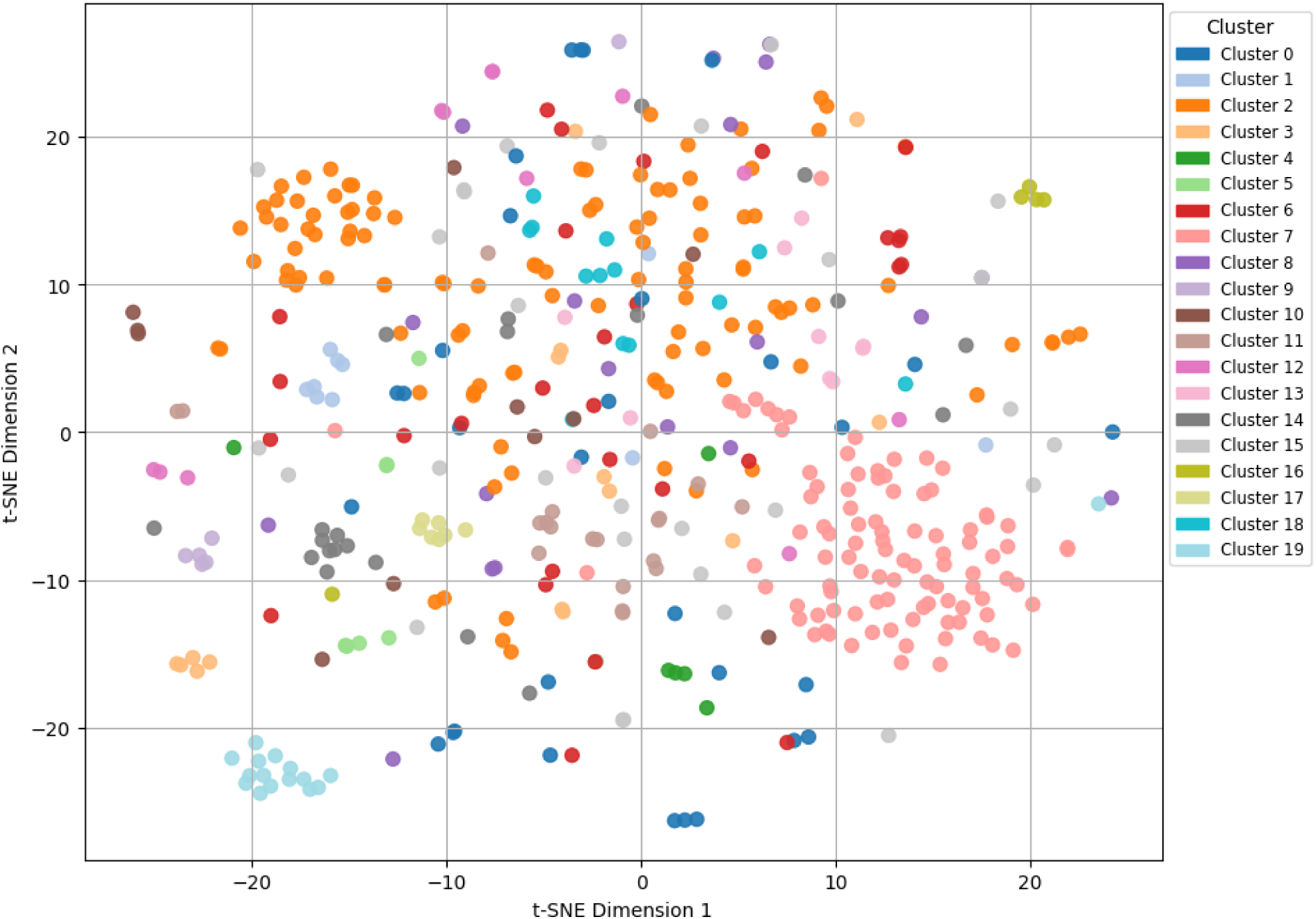
2D t-SNE analysis for the MATGAT-based similarity matrix on the binding-site sequence. The proteins are coloured per clusters as derived by K-medoids PAM clustering algorithm basing to their TMscore.

Figure 5 clearly evidences that some clusters are closely grouped in the plot thus confirming a marked similarity in the binding sites of the proteins belonging to these clusters, while other clusters are spread in the plot or reveal an in-between behaviour. While avoiding a systematic analysis, some clusters deserve attention for their exemplificative behaviour and for the biological role of the corresponding targets. For example, cluster 7 is a clear example of well grouped (pink) points: it comprises 76 kinases which are indeed characterized by highly conserved binding sites. Similar behaviour is shown by cluster 19 (azure points) which contains 15 factors involved in proteolytic cascades. While comprising 16 nuclear hormone receptors, the brown points of cluster 11 are not so closely grouped thus highlighting a certain degree of diversity between their binding pockets. Finally, the cluster 2 shows an intermediate behaviour since there is a clear group of close orange points at top left of the plot, while the remaining points are spread in the plot (especially along the first t-SNE dimension). This behaviour can be explained by considering that cluster 2 includes 107 GPCRs and suggests that some GPCRs exhibit a marked similarity between their binding sites, while many other GPCRs are characterized by less conserved pockets.

The heatmap of Figure 6 shows the two triangular matrices as obtained by analysing the sequence (based on the MatGAT analysis on the entire protein, upper triangle) and the structure (based on TMscore values, lower triangle) similarity between pairs of proteins. Although the density of this map prevents the analysis of specific similarity values, the overall comparison of the two triangular matrices reveals that the structural similarity between the 477 selected targets is higher than the sequence similarity. This result agrees with several published studies (e.g. see ref [40]), which report that protein structure is not markedly affected by residue substitutions, which preserve the sequence hydropathic profile, and proteins with a sequence identity higher than 35–40% usually show a similar folding. Such an observation can also be explained by considering that the proteins can assume a defined number of folding topologies and thus they can share rather similar structures while having a reduced degree of sequence similarity.

**Figure 6.**
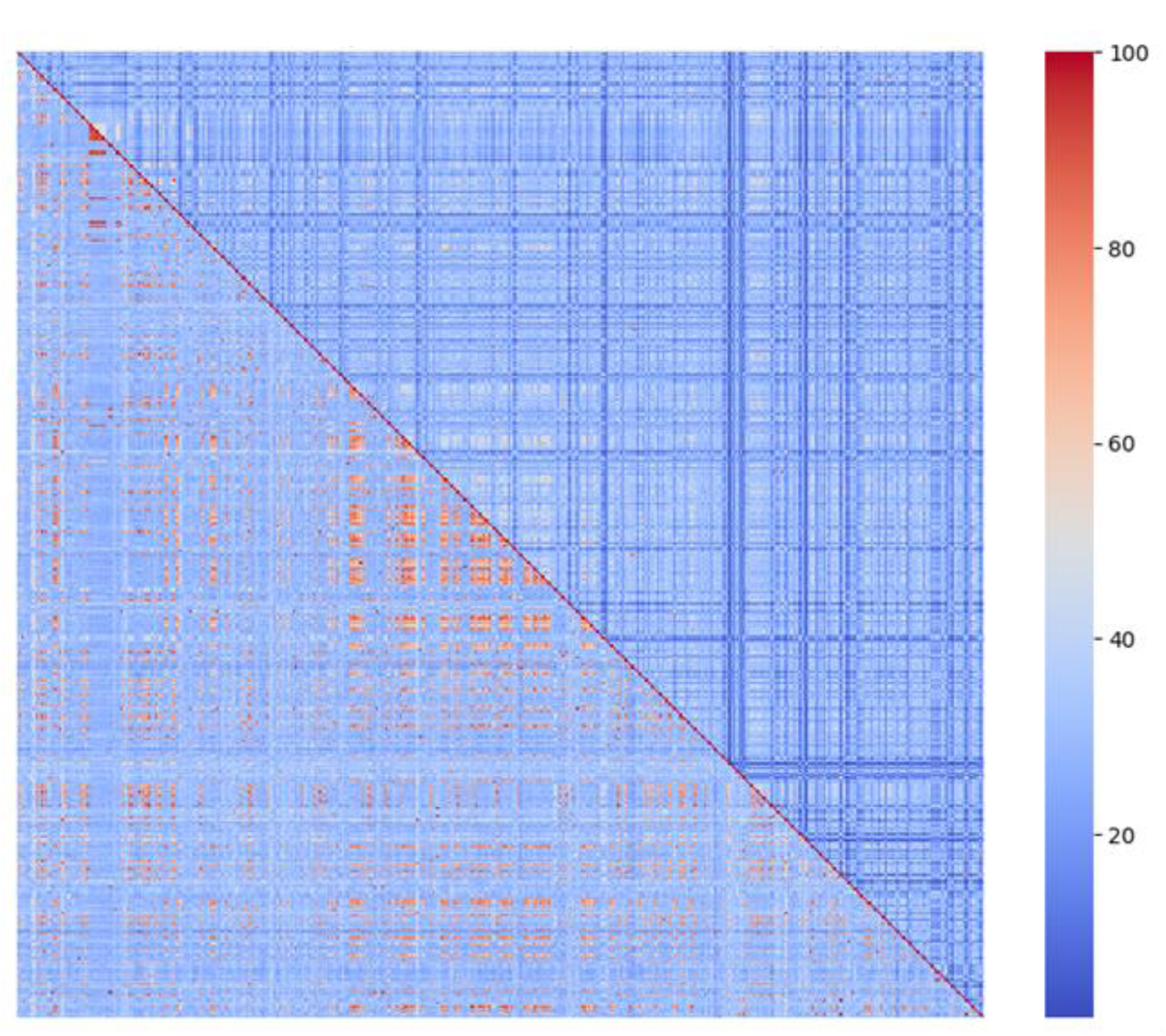
Heatmap for the sequence (upper triangle) and structure (lower triangle) similarity for all the 477 proteins.

The structural similarity is often reflected in the function similarity, and this also depends on the observation that the similarity in the binding sites is, on average, higher than in the overall sequence as shown by heatmap of Figure A2, which compares the sequence similarity for entire protein (lower triangle) and for the binding site only (upper triangle). This confirms that the proteins can share similar binding pockets (and reasonably similar functions) while having quite different overall sequences.

### Overall analysis

Inspired by the previous analyses of both ligands and targets, a combined analysis was performed to evaluate if similar targets bind similar ligands or, in other words, to explore the correlation between target and ligand similarity. Thus, a PCA analysis involved the 477 sets of active ligands which were characterized by calculating the average values for 26 VEGA ZZ-based physicochemical descriptors, as already done for Figure 1.

Figure 7 shows the two resulting principal components in which the points are coloured by protein clusters based on their TMscore as already discussed for Figure 5. While not showing a clear grouping of the points of the same cluster, Figure 7 shows that the points of the five analysed clusters (with more than 25 proteins) are spread along the PC1 axis, while revealing rather constant PC2 values. This behaviour is particularly evident for cluster 7 (red points) which is composed by 76 kinases, while being less clear for cluster 2, which comprises 107 GPCRs. In agreement with what was seen in Figure 1, the features mostly contributing to the PC1 values (Table A7) are related to molecular size, while those contributing to the PC2 values (Table A8) are focused on molecular polarity. Hence, the PCA plot in Figure 7 suggests that the ligands of similar targets share similar polarity while possessing different molecular shapes. Similar results (despite a bit less clear) are obtained when clustering the proteins based on the MatGAT analysis on the entire protein or on the binding site only (data not shown).

**Figure 7.**
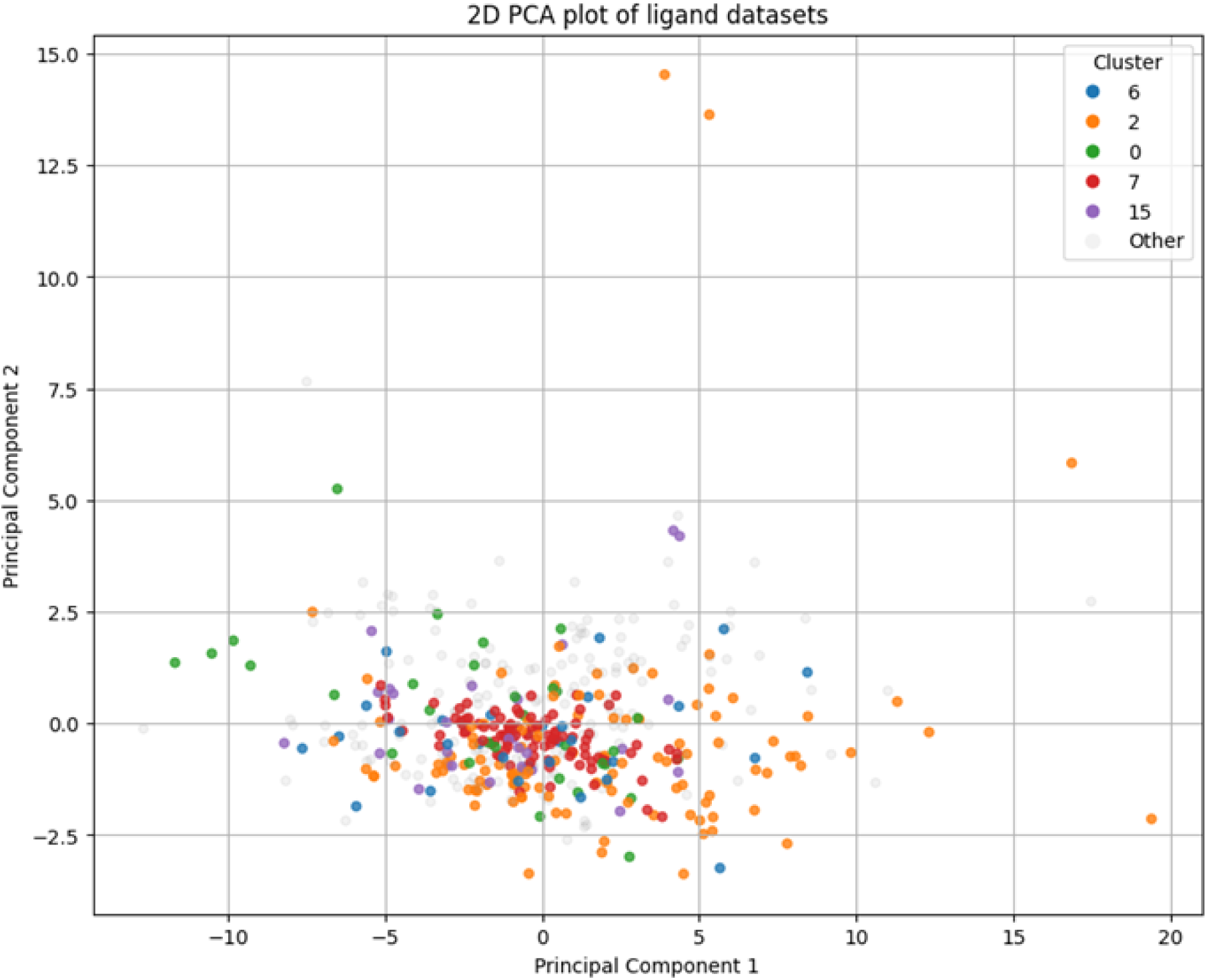
PCA analysis of 26 VEGA ZZ-based physicochemical descriptors for the 477 active datasets. The points are coloured by protein clusters according to their TMscore. Only clusters including more than 25 proteins are displayed.

To extend the analysis to all collected proteins, the triangular similarity matrices as shown in the previously discussed heatmaps (see Figure 6 and A2) and focused on protein structures were correlated with the corresponding triangular matrices as obtained by computing pairwise similarity distances for the active ligands. In detail, for each set of active molecules and for each set of descriptors the average values of the selected descriptors were utilized to calculate the Euclidean distances (in N-dimensional space, where N is the number of descriptors in the considered set) between pairs of targets. The so-obtained distance values were further normalized in the range 0-100 and converted into ligand similarity values by applying the equation: similarity = 100 – normalized distance.

Table 2 reports the Pearson coefficients (r) as derived when correlating the ligand similarity matrices based on VEGA ZZ, MOPAC and RDKit descriptors plus Morgan fingerprints with the protein similarity values based on TM and MATGAT scores. While showing rather low values, the compiled Pearson coefficients allow for some relevant considerations. Concerning the ligand analyses, the Morgan fingerprints provide the best correlations thus highlighting the key role of structural similarity compared to physicochemical or stereo-electronic similarity. Concerning the protein structure, the similarity of the binding pockets yields the best correlations thus emphasizing the relevance of the binding cavities in determining the protein features. To summarize what reported in Table 2, one may conclude that there is a weak but not negligible correlation between ligand and target similarity or, stated differently, that similar binding sites tend to recognize rather similar ligands.

**Table 2.** Pearson coefficients for the correlations between the matrices for protein similarity (involving the protein structure, TMscore, as well as the sequence of the entire protein and of the sole binding site) and ligand similarity based on four sets of fingerprints and descriptors.

| Descriptors | TM-score | MATGAT score entire protein | MATGAT score binding site |
| --- | --- | --- | --- |
| VEGA ZZ | 0.099 | 0.052 | 0.162 |
| Morgan | 0.111 | 0.158 | 0.224 |
| RDKit | 0.131 | 0.094 | 0.206 |
| MOPAC | 0.117 | 0.061 | 0.149 |

### Polypharmacological profile

The collected ligands and the related biological data also allow a detailed analysis of their polypharmacological profile since many molecules are characterized by activity values experimentally determined for more than one target. In detail, 70832 molecules are described by biological data for one only target, while 8106 molecules possess biological data for more than one target, that can be mapped into 24568 data points. The 8106 molecules with multiple biological data can be subdivided into: (1) 2353 ligands active on all tested targets; (2) 2806 ligands inactive on all tested targets; (3) 2947 ligands active on some targets and inactive on other targets.

When focusing on molecules active on all tested targets, Table 3 shows that a vast majority (about 80%) is active on two targets followed by molecules active on three targets (13.5%) even though there are nine molecules active on more than 10 targets with two cases being active on 26 and 29 targets. The analysis of the biological classes of the targets for the 1871 molecules active on two targets (see Table A9) reveals that the two targets generally belong to the same biological class. Indeed, only 34 molecules (1.8%) are active on one enzyme and one receptor despite the relevance of these two classes among the collected targets. Similar homogeneous results are obtained when analysing the 318 compounds active on three targets (data not shown).

**Table 3.**
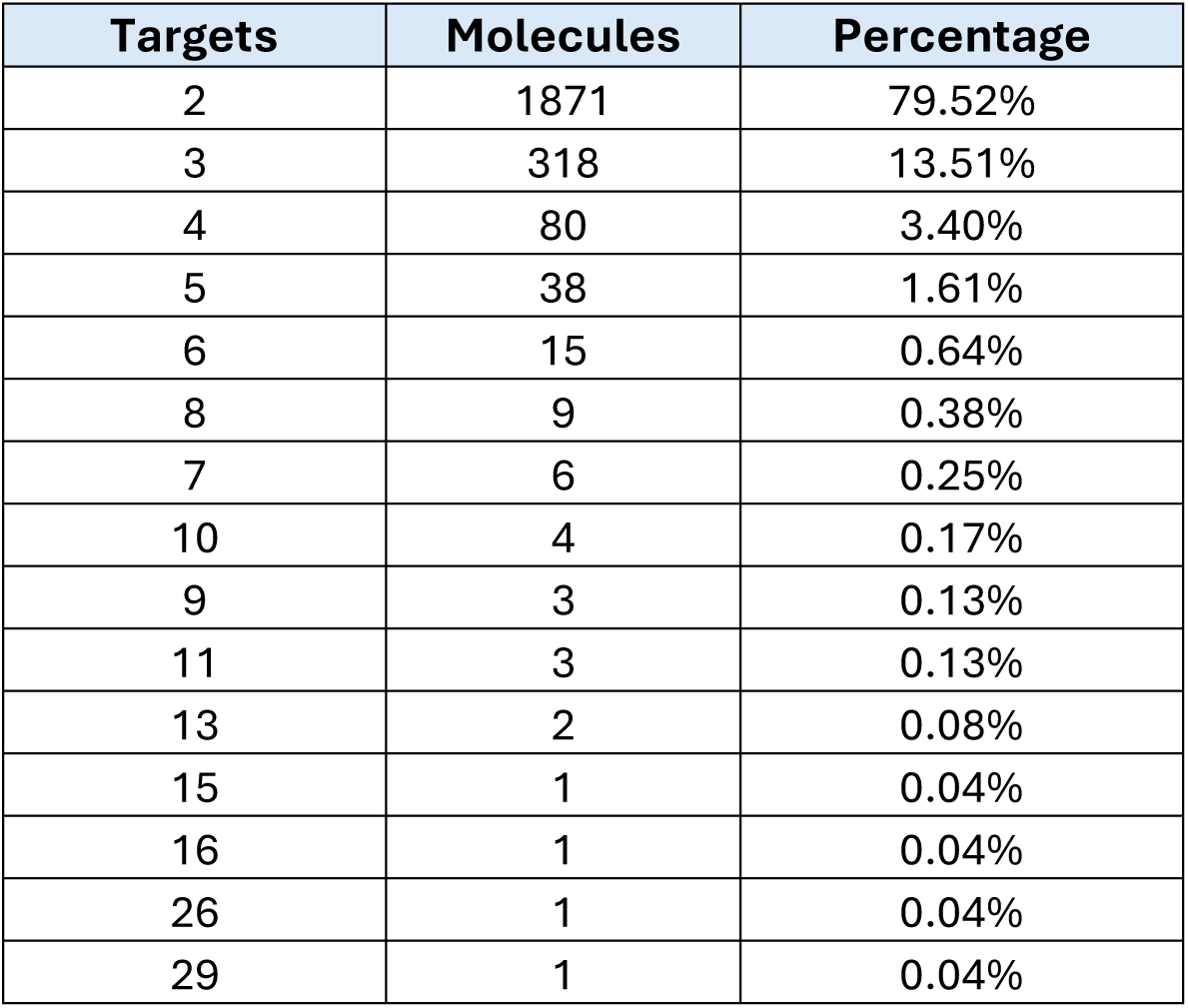
Distribution of the 2353 ligands which are active on more than one target.

The analysis of molecules inactive on all tested targets shows similar results (see Table A10) with molecules inactive on two or three targets which correspond to about 90%. Also here, there are molecules (10) which are inactive on more than 15 targets, with one molecule being inactive on 27 targets. The analysis of the involved targets for the 2209 ligands inactive on two targets confirms that the targets generally belong to the same class (data not shown).

Lastly, the analysis of molecules with mixed active and inactive profile (Table A11) shows that about one half is active on one target and inactive on one target, while the two combinations of molecules active on one target and inactive on two targets and vice versa correspond to about 18%. Table A10 also highlights the high number of retrieved combinations (163), with 30 and 8 molecules characterized by more than 20 and 30 biological data, respectively, with a representative case which is active on 39 targets and inactive on one target. The analysis of the involved targets for the 1476 ligands with two biological data confirms that the targets belong to the same biological classes in most cases, even though the combination one enzyme plus one receptor here corresponds to about 5% with 67 cases (data not shown). Similar conclusions can be drawn when analysing the 534 molecules with three biological data.

### Description of the DELTA website

All the collected and prepared data are available for download from the website delta.unimi.it. Figure 8 shows the structure of the webpage dedicated to the target structures (on the top) as well as an example of webpage reporting the collected ligands for a given target (on the bottom). As evidenced in Figure 8 (top panel) and for each target, the table compiles the name, the UniProt AC and ID codes, the biological class, the PDB ID for experimentally resolved structures and if that target is of a marketed drug or in clinical trial. For each target, the prepared protein structure is available for download in mol2 format and can be viewed by a visualization webpage powered by the WebGL-based visualization library [41]. For each target, the table also includes the link to the webpage for the corresponding ligands. As depicted in Figure 8, the target webpage also comprises search tools by which the target(s) can be variously searched, and similar tools are also implemented in the visualization webpage.

**Figure 8.**
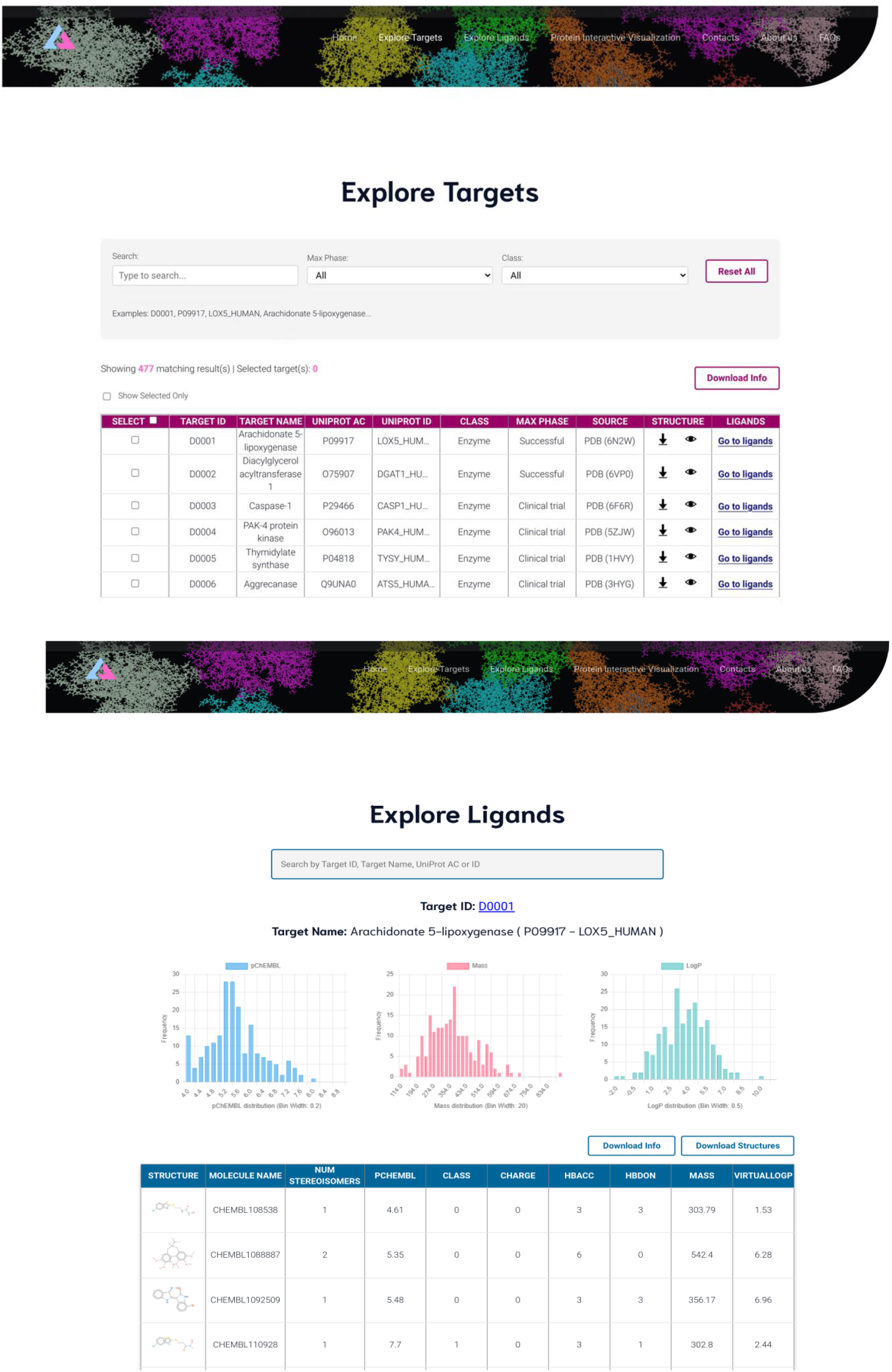
DELTA webpage for the targets (top panel) and an example of webpage for the ligands (bottom panel).

For each target, the ligands’ webpage (see Figure 8, bottom panel) includes a table compiling the 200 collected ligands. For each ligand, the table includes the 2D structure, the ChEMBL name, the number of considered stereoisomers, the pChEMBL, the class (active/inactive) plus a set of descriptors such as molecular mass, charge, number of H-bond groups and virtual logP value. To better characterize the collected ligands, the webpage also includes three plots showing the distribution of pChEMBL values, mass and lipophilicity. Each webpage allows downloading the prepared 3D structures of the ligands (in mol2 format) as well as a CSV file including for each ligand the VEGA ZZ-based physicochemical descriptors plus the stereo-electronic features as computed by PM7 semi-empirical calculations. Although the DELTA website was designed to enable specific downloads, it allows the download of a CSV file comprising all the collected ligands and including the SMILES strings along with a set of relevant descriptors.

## Conclusions

The here proposed DELTA resource provides curated and balanced ligands’ datasets for about 500 therapeutically relevant targets plus the optimized structure of the corresponding target. The included data are prepared to develop AI-based classification models by using both ligand- and structure-based approaches. Along with this primary application, the DELTA resource can find many other applications. As an example, the DELTA database can be used to generate enhanced datasets for virtual screening validations by dispersing the 100 active ligands for a given target with a lot of presumably inactive ligands suitably chosen among all the other ligands. More in general, the DELTA ligands can represent a valuable dataset for virtual screening campaigns by considering all ligands, all active ligands, or all ligands with pChEMBL greater than a defined threshold. Furthermore, the DELTA resource is a clearly flexible and easily expandable system by adding new targets with relative ligands’ datasets or by increasing the number of collected ligands per target (when available). As described above, all the prepared structures (targets plus ligands) are available for download at delta.unimi.it which is structured for specific download. Nevertheless, the entire DELTA resource can be obtained upon request to the Authors (sending an email to).

## Acknowledgement

This work was supported by the Project Multilayered Urban Sustainability Action (MUSA), funded by the European Union–NextGenerationEU, under the National Recovery and Resilience Plan (NRRP) Mission Four Component Two Investment Line 1.5.

## Supporting Information

**Figure A1.**
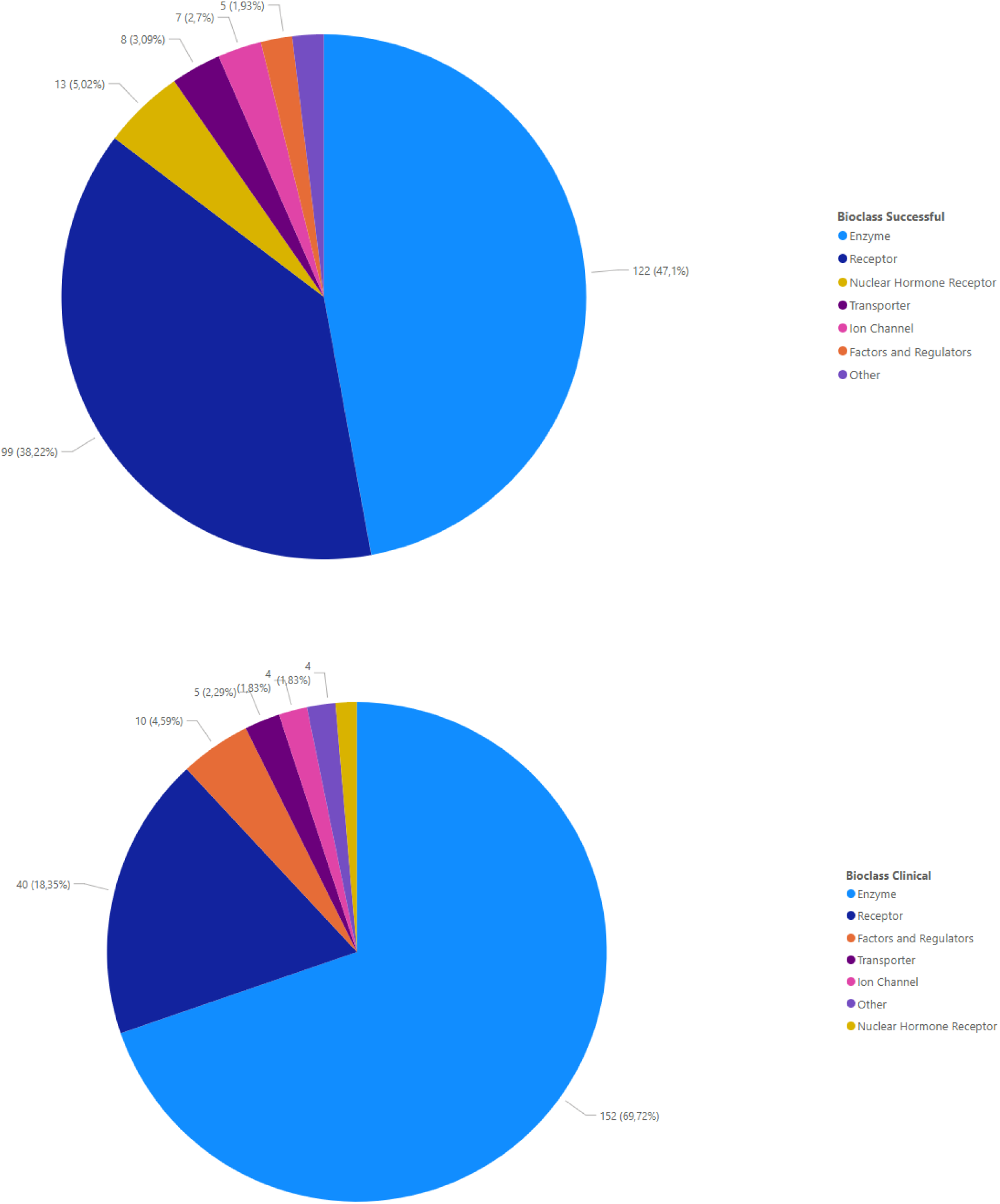
Classification of the targets for marketed drugs (on the top) and for drugs in clinical trials (on the bottom) based on their biological class.

**Figure A2.**
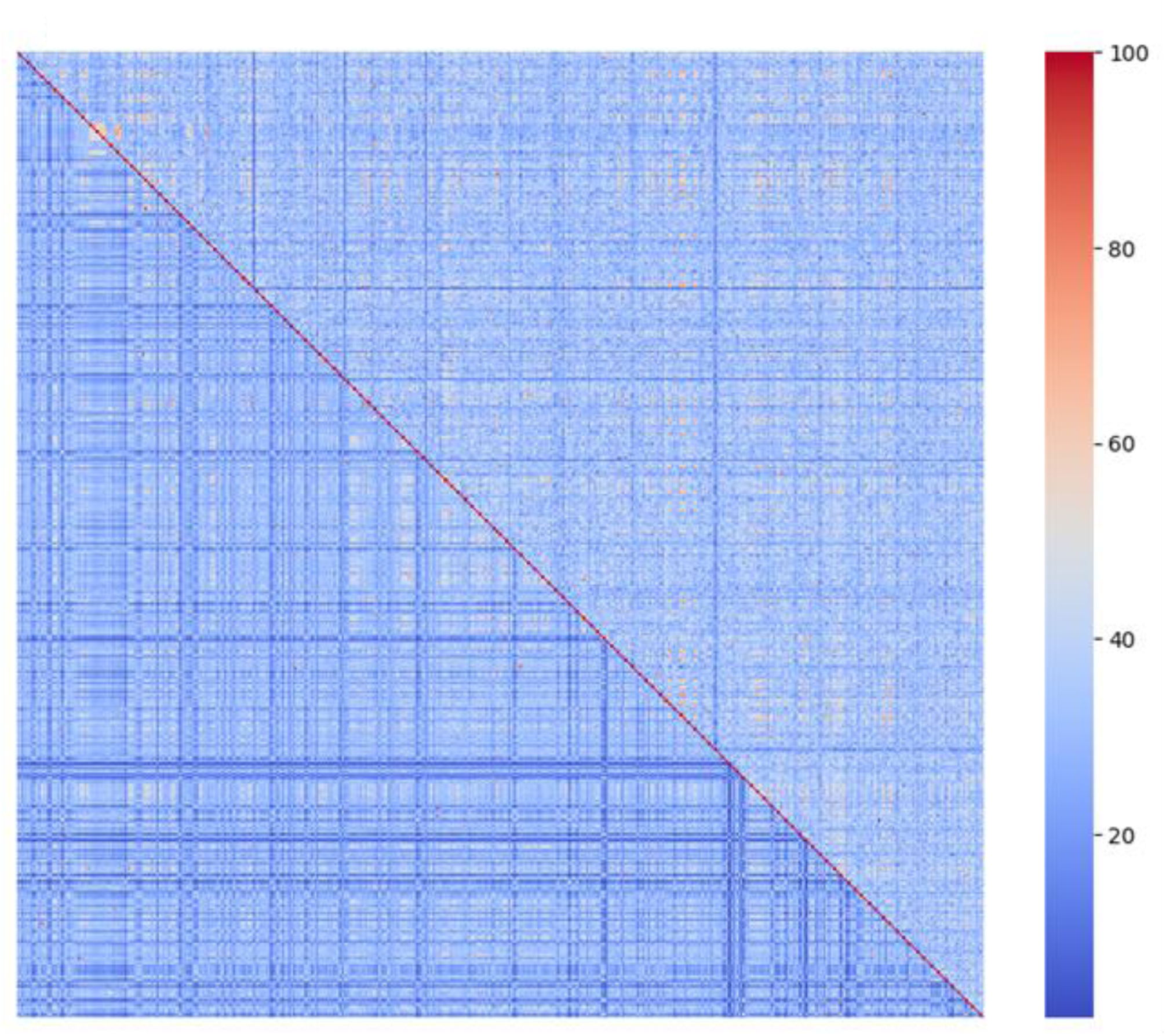
Heatmap for the sequence similarity for the entire protein (lower triangle) and for the binding site only (upper triangle) for all the 477 proteins.

---

**Table A1.** PCA loadings on the first two components (PC1, Table A1 and PC2, Table A2) as depicted in Figure 1 and based on physicochemical descriptors.

| VEGA ZZ descriptor | Loading on PC1 |
| --- | --- |
| Sav | 0.995674699 |
| Volume | 0.994741956 |
| Surface | 0.993706453 |
| Sdiam | 0.992901835 |
| Vdiam | 0.990750592 |
| HeavyAtoms | 0.988764325 |
| Sas | 0.984169945 |
| Ovality | 0.982099107 |
| Mass | 0.980784683 |
| Bonds | 0.978583485 |
| Atoms | 0.976163495 |
| Angles | 0.953873659 |
| Torsions | 0.93300792 |
| FlexTorsions | 0.861148436 |
| Gyrrad | 0.825935774 |
| Rings | 0.633543642 |
| Impropers | 0.582257151 |
| Psa | 0.563342428 |
| HbAcc | 0.519199458 |
| ChiralAtms | 0.429447985 |
| HbDon | 0.369252133 |
| Dipole | 0.300523981 |
| VirtualLogP | 0.288548541 |
| Charge | 0.020379957 |
| Lipole | -0.016042839 |
| EzBnds | -0.078989782 |

**Table A2.**
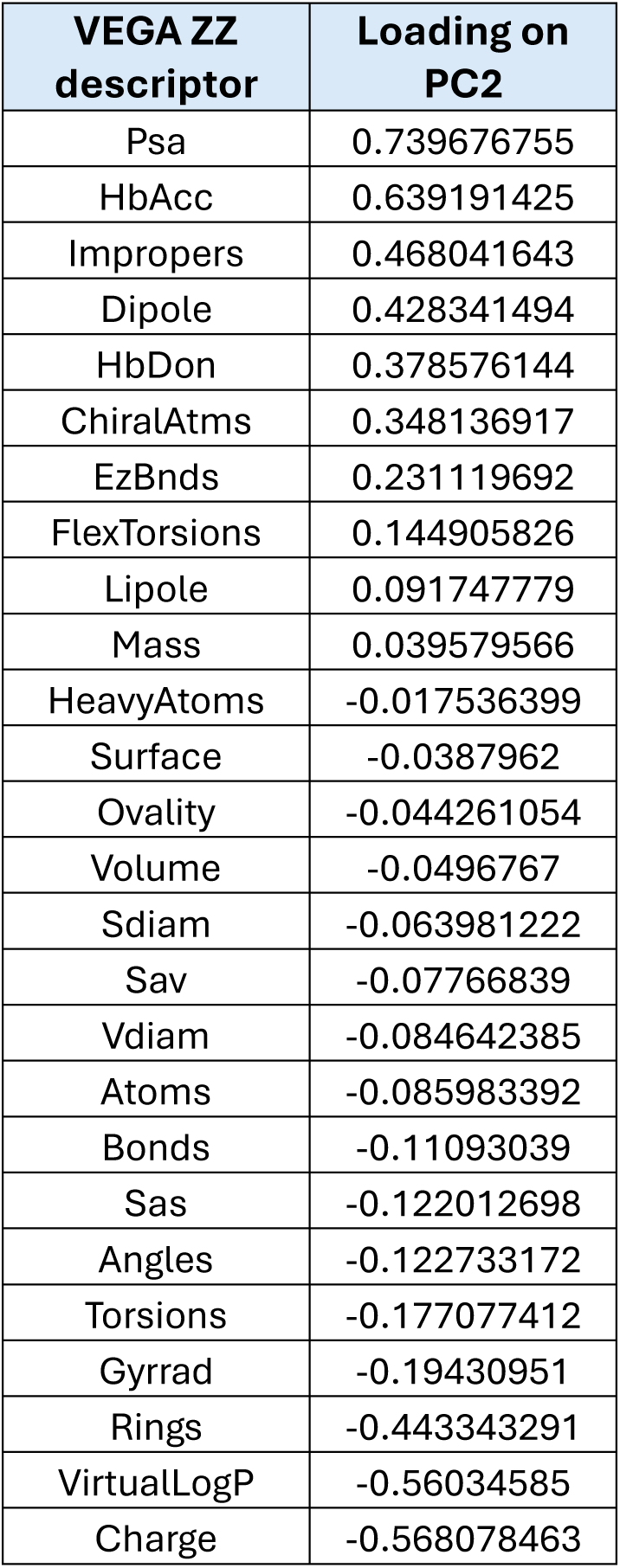
PCA loadings on the first two components (PC1, Table A1 and PC2, Table A2) as depicted in Figure 1 and based on physicochemical descriptors.

| VEGA ZZ descriptor | Loading on PC2 |
| --- | --- |
| Psa | 0.739676755 |
| HbAcc | 0.639191425 |
| Impropers | 0.468041643 |
| Dipole | 0.428341494 |
| HbDon | 0.378576144 |
| ChiralAtms | 0.348136917 |
| EzBnds | 0.231119692 |
| FlexTorsions | 0.144905826 |
| Lipole | 0.091747779 |
| Mass | 0.039579566 |
| HeavyAtoms | -0.017536399 |
| Surface | -0.0387962 |
| Ovality | -0.044261054 |
| Volume | -0.0496767 |
| Sdiam | -0.063981222 |
| Sav | -0.07766839 |
| Vdiam | -0.084642385 |
| Atoms | -0.085983392 |
| Bonds | -0.11093039 |
| Sas | -0.122012698 |
| Angles | -0.122733172 |
| Torsions | -0.177077412 |
| Gyrrad | -0.19430951 |
| Rings | -0.443343291 |
| VirtualLogP | -0.56034585 |
| Charge | -0.568078463 |

**Table A3.** Explained variance with the first two principal components using different sets of descriptors plus their combinations.

| Descriptors | Explained variance |
| --- | --- |
| VEGA ZZ | 0.71 |
| MOPAC | 0.62 |
| RDKit | 0.43 |
| VEGA ZZ + MOPAC | 0.63 |
| VEGA ZZ + RDKit | 0.47 |
| RDKit+ MOPAC | 0.45 |
| VEGA ZZ + MOPAC + RDKit | 0.48 |

**Table A4.**
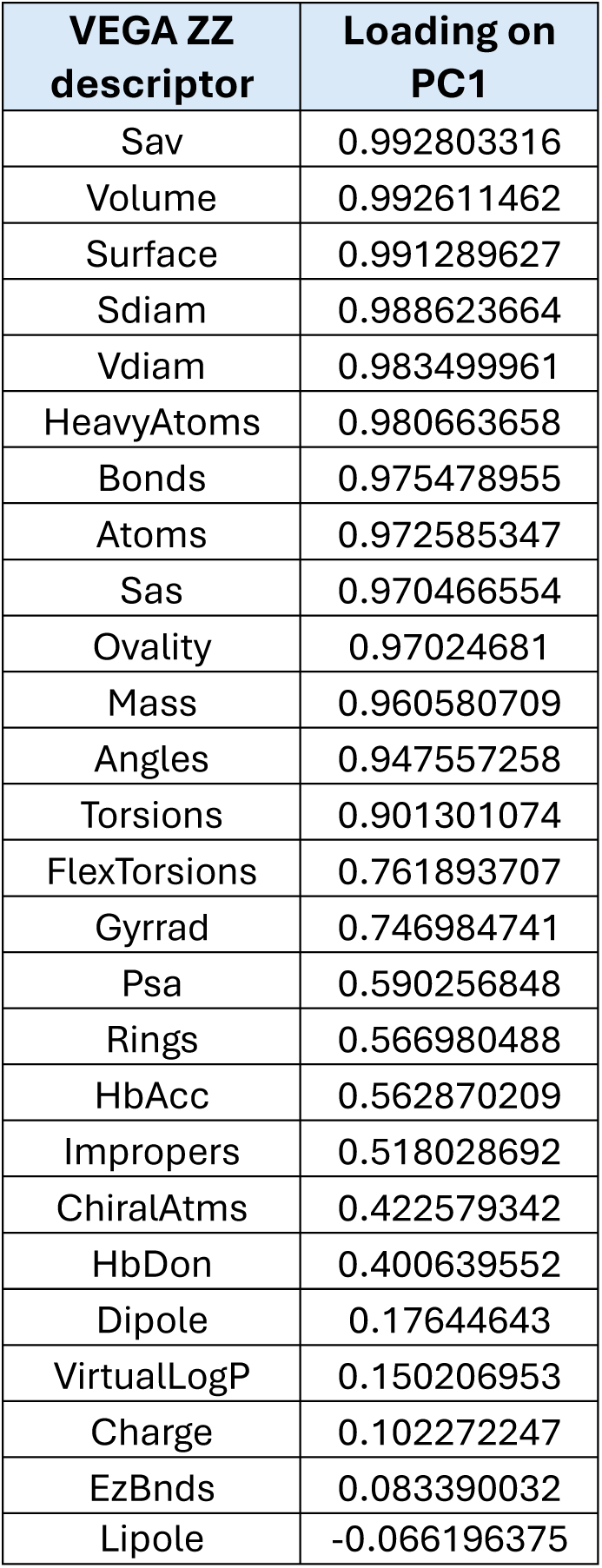
PCA loadings on the first two components (PC1, Table A4 and PC2, Table A5) as depicted in Figure 2 and based on physicochemical descriptors.

| VEGA ZZ descriptor | Loading on PC1 |
| --- | --- |
| Sav | 0.992803316 |
| Volume | 0.992611462 |
| Surface | 0.991289627 |
| Sdiam | 0.988623664 |
| Vdiam | 0.983499961 |
| HeavyAtoms | 0.980663658 |
| Bonds | 0.975478955 |
| Atoms | 0.972585347 |
| Sas | 0.970466554 |
| Ovality | 0.97024681 |
| Mass | 0.960580709 |
| Angles | 0.947557258 |
| Torsions | 0.901301074 |
| FlexTorsions | 0.761893707 |
| Gyrrad | 0.746984741 |
| Psa | 0.590256848 |
| Rings | 0.566980488 |
| HbAcc | 0.562870209 |
| Impropers | 0.518028692 |
| ChiralAtms | 0.422579342 |
| HbDon | 0.400639552 |
| Dipole | 0.17644643 |
| VirtualLogP | 0.150206953 |
| Charge | 0.102272247 |
| EzBnds | 0.083390032 |
| Lipole | -0.066196375 |

**Table A5.** PCA loadings on the first two components (PC1, Table A4 and PC2, Table A5) as depicted in Figure 2 and based on physicochemical descriptors.

| VEGA ZZ descriptor | Loading on PC2 |
| --- | --- |
| VirtualLogP | 0.773618418 |
| Rings | 0.420365412 |
| Gyrrad | 0.232098385 |
| Torsions | 0.18075375 |
| Sas | 0.139382415 |
| Charge | 0.118657463 |
| Vdiam | 0.105421827 |
| Lipole | 0.090237185 |
| Sav | 0.083331711 |
| Sdiam | 0.073810124 |
| Bonds | 0.061542517 |
| Angles | 0.057747864 |
| Volume | 0.049800244 |
| HeavyAtoms | 0.048422883 |
| Ovality | 0.045379621 |
| Atoms | 0.03288037 |
| Surface | 0.032583227 |
| Mass | 0.017370086 |
| FlexTorsions | -0.12603964 |
| EzBnds | -0.150422084 |
| Dipole | -0.174891377 |
| ChiralAtms | -0.338923938 |
| HbAcc | -0.43434241 |
| Impropers | -0.478278347 |
| HbDon | -0.647200088 |
| Psa | -0.667707633 |

**Table A6.**
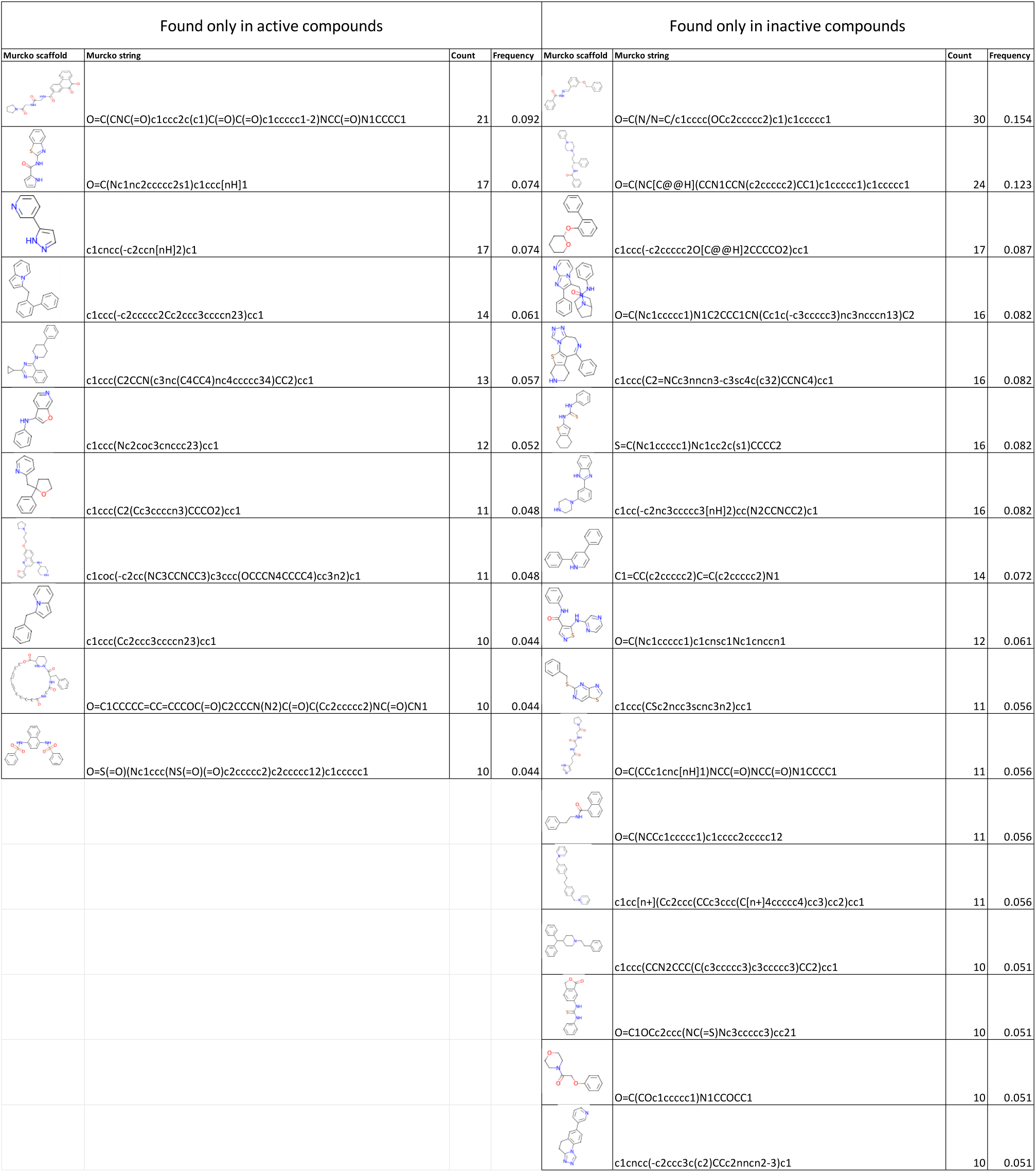
Scaffolds which are present at least 10 times only in the active (on the left) or only in the inactive (on the right) compounds.

**Table A7.** PCA loadings on the first two components (PC1, Table A4 and PC2, Table A5) as depicted in Figure 7 and based on physicochemical descriptors.

| VEGA ZZ descriptor | Loading on PC1 |
| --- | --- |
| Sav | 0.996134 |
| Volume | 0.99508 |
| Surface | 0.994192 |
| Sdiam | 0.993553 |
| Vdiam | 0.991296 |
| HeavyAtoms | 0.988664 |
| Sas | 0.983439 |
| Ovality | 0.983124 |
| Mass | 0.980429 |
| Bonds | 0.978756 |
| Atoms | 0.976137 |
| Angles | 0.954353 |
| Torsions | 0.930237 |
| FlexTorsions | 0.86104 |
| Gyrrad | 0.806364 |
| Rings | 0.616592 |
| Impropers | 0.614274 |
| Psa | 0.56183 |
| HbAcc | 0.530861 |
| ChiralAtms | 0.44276 |
| HbDon | 0.366634 |
| VirtualLogP | 0.293896 |
| Dipole | 0.244468 |
| Charge | 0.0118461 |
| Lipole | -0.0516574 |
| EzBnds | -0.116769 |

**Table A8.** PCA loadings on the first two components (PC1, Table A4 and PC2, Table A5) as depicted in Figure 7 and based on physicochemical descriptors.

| VEGA ZZ descriptor | Loading on PC2 |
| --- | --- |
| Psa | 0.742377 |
| HbAcc | 0.648112 |
| Impropers | 0.48418 |
| Dipole | 0.417795 |
| HbDon | 0.344092 |
| ChiralAtms | 0.330884 |
| EzBnds | 0.324377 |
| FlexTorsions | 0.158196 |
| Lipole | 0.0699587 |
| Mass | 0.0480268 |
| HeavyAtoms | -0.0149288 |
| Surface | -0.0355387 |
| Ovality | -0.0376799 |
| Volume | -0.0476159 |
| Sdiam | -0.0601258 |
| Sav | -0.0776638 |
| Vdiam | -0.082112 |
| Atoms | -0.0874113 |
| Bonds | -0.112099 |
| Angles | -0.123281 |
| Sas | -0.127026 |
| Torsions | -0.181365 |
| Gyrrad | -0.211409 |
| Rings | -0.445709 |
| VirtualLogP | -0.556936 |
| Charge | -0.578315 |

**Table A9.**
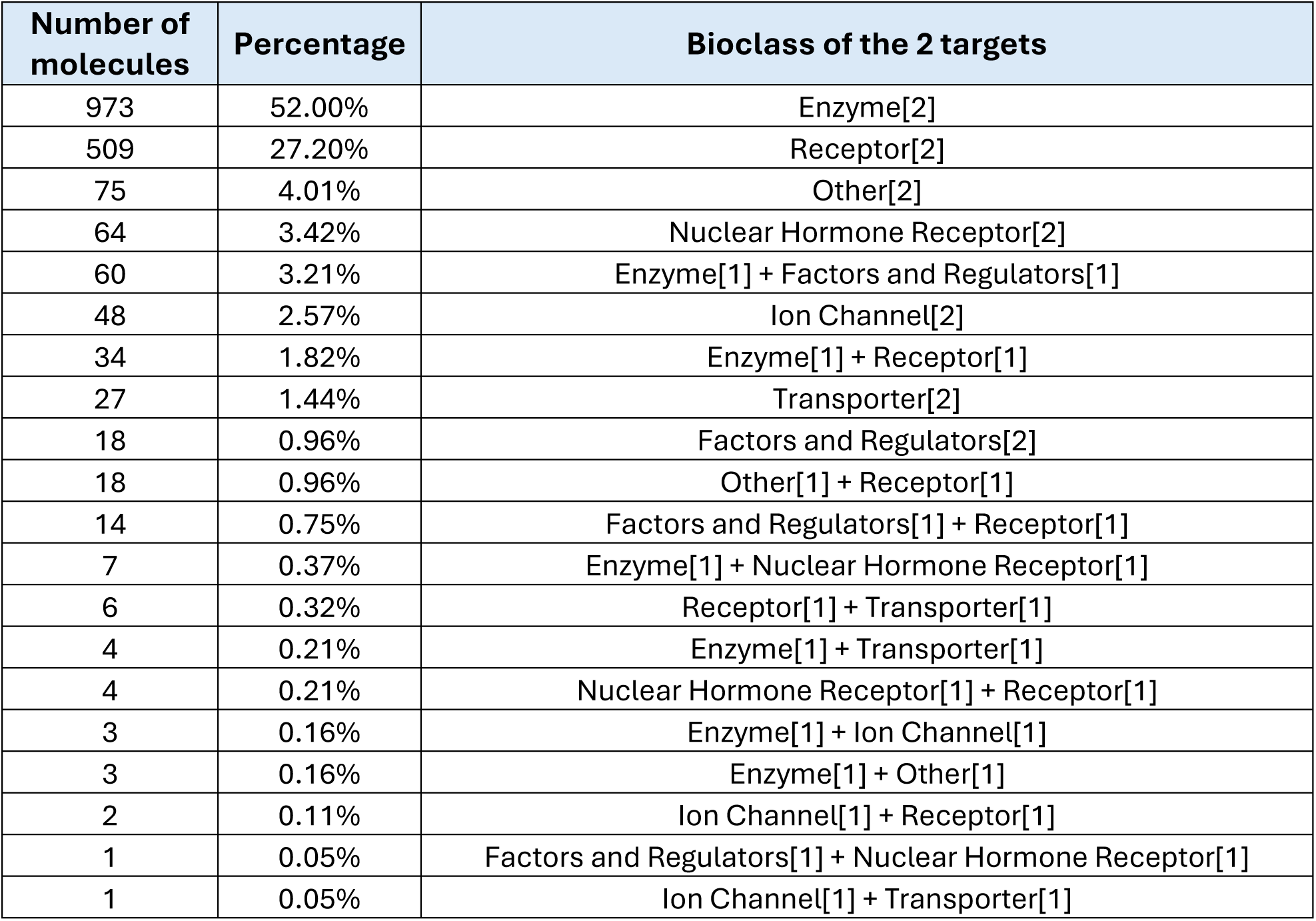
Distribution of the biological classes of the targets for the 1871 molecules which are active on two targets.

**Table A10.** Distribution of the 2806 ligands which are inactive on more than one target.

| Targets | Molecules | Percentage |
| --- | --- | --- |
| 2 | 2209 | 78.72% |
| 3 | 375 | 13.36% |
| 4 | 107 | 3.81% |
| 5 | 57 | 2.03% |
| 6 | 12 | 0.43% |
| 7 | 8 | 0.29% |
| 9 | 6 | 0.21% |
| 11 | 6 | 0.21% |
| 8 | 5 | 0.18% |
| 10 | 4 | 0.14% |
| 15 | 3 | 0.11% |
| 14 | 2 | 0.07% |
| 16 | 2 | 0.07% |
| 17 | 2 | 0.07% |
| 21 | 2 | 0.07% |
| 12 | 1 | 0.04% |
| 13 | 1 | 0.04% |
| 18 | 1 | 0.04% |
| 19 | 1 | 0.04% |
| 20 | 1 | 0.04% |
| 27 | 1 | 0.04% |

**Table A11.** Distribution of the 2947 ligands which are active on some targets and inactive on other targets.

| Targets on which molecule is active | Targets on which molecule is inactive | Molecules | Percentage |
| --- | --- | --- | --- |
| 1 | 1 | 1476 | 50.08% |
| 1 | 2 | 306 | 10.38% |
| 2 | 1 | 228 | 7.74% |
| 1 | 3 | 92 | 3.12% |
| 2 | 2 | 90 | 3.05% |
| 3 | 1 | 87 | 2.95% |
| 1 | 4 | 43 | 1.46% |
| 2 | 3 | 38 | 1.29% |
| 4 | 1 | 37 | 1.26% |
| 3 | 2 | 35 | 1.19% |
| 2 | 4 | 25 | 0.85% |
| 5 | 1 | 23 | 0.78% |
| 1 | 5 | 19 | 0.64% |
| 5 | 2 | 17 | 0.58% |
| 3 | 3 | 17 | 0.58% |
| 5 | 3 | 15 | 0.51% |
| 4 | 2 | 15 | 0.51% |
| 4 | 3 | 14 | 0.48% |
| 3 | 5 | 13 | 0.44% |
| 7 | 2 | 10 | 0.34% |
| 9 | 2 | 9 | 0.31% |
| 3 | 4 | 9 | 0.31% |
| 2 | 5 | 9 | 0.31% |
| 1 | 6 | 9 | 0.31% |
| 4 | 5 | 8 | 0.27% |
| 4 | 4 | 8 | 0.27% |
| 6 | 1 | 8 | 0.27% |
| 10 | 1 | 7 | 0.24% |
| 7 | 3 | 7 | 0.24% |
| 5 | 4 | 7 | 0.24% |
| 2 | 10 | 6 | 0.20% |
| 4 | 8 | 6 | 0.20% |
| 7 | 4 | 6 | 0.20% |
| 6 | 5 | 6 | 0.20% |
| 6 | 4 | 6 | 0.20% |
| 5 | 5 | 6 | 0.20% |
| 4 | 6 | 6 | 0.20% |
| 8 | 1 | 6 | 0.20% |
| 6 | 2 | 6 | 0.20% |
| 1 | 7 | 6 | 0.20% |
| 11 | 2 | 5 | 0.17% |
| 9 | 1 | 5 | 0.17% |
| 8 | 2 | 5 | 0.17% |
| 14 | 2 | 4 | 0.14% |
| 10 | 5 | 4 | 0.14% |
| 10 | 3 | 4 | 0.14% |
| 10 | 2 | 4 | 0.14% |
| 9 | 3 | 4 | 0.14% |
| 8 | 4 | 4 | 0.14% |
| 3 | 8 | 4 | 0.14% |
| 6 | 3 | 4 | 0.14% |
| 2 | 6 | 4 | 0.14% |
| 13 | 4 | 3 | 0.10% |
| 9 | 5 | 3 | 0.10% |
| 9 | 4 | 3 | 0.10% |
| 8 | 5 | 3 | 0.10% |
| 7 | 5 | 3 | 0.10% |
| 6 | 6 | 3 | 0.10% |
| 5 | 6 | 3 | 0.10% |
| 3 | 7 | 3 | 0.10% |
| 1 | 9 | 3 | 0.10% |
| 2 | 7 | 3 | 0.10% |
| 18 | 6 | 2 | 0.07% |
| 17 | 5 | 2 | 0.07% |
| 10 | 10 | 2 | 0.07% |
| 16 | 4 | 2 | 0.07% |
| 13 | 7 | 2 | 0.07% |
| 15 | 3 | 2 | 0.07% |
| 11 | 6 | 2 | 0.07% |
| 7 | 10 | 2 | 0.07% |
| 9 | 7 | 2 | 0.07% |
| 13 | 2 | 2 | 0.07% |
| 12 | 3 | 2 | 0.07% |
| 2 | 13 | 2 | 0.07% |
| 8 | 7 | 2 | 0.07% |
| 7 | 8 | 2 | 0.07% |
| 12 | 2 | 2 | 0.07% |
| 11 | 3 | 2 | 0.07% |
| 7 | 7 | 2 | 0.07% |
| 5 | 9 | 2 | 0.07% |
| 2 | 11 | 2 | 0.07% |
| 1 | 12 | 2 | 0.07% |
| 8 | 3 | 2 | 0.07% |
| 4 | 7 | 2 | 0.07% |
| 2 | 8 | 2 | 0.07% |
| 3 | 6 | 2 | 0.07% |
| 7 | 1 | 2 | 0.07% |
| 34 | 7 | 1 | 0.03% |
| 39 | 1 | 1 | 0.03% |
| 6 | 33 | 1 | 0.03% |
| 30 | 2 | 1 | 0.03% |
| 26 | 6 | 1 | 0.03% |
| 23 | 9 | 1 | 0.03% |
| 28 | 3 | 1 | 0.03% |
| 8 | 23 | 1 | 0.03% |
| 27 | 2 | 1 | 0.03% |
| 25 | 3 | 1 | 0.03% |
| 23 | 5 | 1 | 0.03% |
| 20 | 8 | 1 | 0.03% |
| 23 | 4 | 1 | 0.03% |
| 24 | 2 | 1 | 0.03% |
| 20 | 6 | 1 | 0.03% |
| 8 | 18 | 1 | 0.03% |
| 16 | 9 | 1 | 0.03% |
| 9 | 16 | 1 | 0.03% |
| 8 | 16 | 1 | 0.03% |
| 12 | 11 | 1 | 0.03% |
| 11 | 12 | 1 | 0.03% |
| 15 | 8 | 1 | 0.03% |
| 1 | 22 | 1 | 0.03% |
| 12 | 10 | 1 | 0.03% |
| 18 | 4 | 1 | 0.03% |
| 19 | 2 | 1 | 0.03% |
| 6 | 15 | 1 | 0.03% |
| 1 | 20 | 1 | 0.03% |
| 19 | 1 | 1 | 0.03% |
| 18 | 2 | 1 | 0.03% |
| 15 | 5 | 1 | 0.03% |
| 14 | 6 | 1 | 0.03% |
| 11 | 9 | 1 | 0.03% |
| 9 | 11 | 1 | 0.03% |
| 18 | 1 | 1 | 0.03% |
| 17 | 2 | 1 | 0.03% |
| 15 | 4 | 1 | 0.03% |
| 10 | 9 | 1 | 0.03% |
| 4 | 15 | 1 | 0.03% |
| 1 | 18 | 1 | 0.03% |
| 17 | 1 | 1 | 0.03% |
| 14 | 4 | 1 | 0.03% |
| 13 | 5 | 1 | 0.03% |
| 8 | 10 | 1 | 0.03% |
| 7 | 11 | 1 | 0.03% |
| 1 | 17 | 1 | 0.03% |
| 16 | 1 | 1 | 0.03% |
| 15 | 2 | 1 | 0.03% |
| 12 | 5 | 1 | 0.03% |
| 10 | 7 | 1 | 0.03% |
| 3 | 14 | 1 | 0.03% |
| 2 | 15 | 1 | 0.03% |
| 15 | 1 | 1 | 0.03% |
| 13 | 3 | 1 | 0.03% |
| 11 | 5 | 1 | 0.03% |
| 10 | 6 | 1 | 0.03% |
| 8 | 8 | 1 | 0.03% |
| 14 | 1 | 1 | 0.03% |
| 5 | 10 | 1 | 0.03% |
| 1 | 14 | 1 | 0.03% |
| 9 | 6 | 1 | 0.03% |
| 6 | 9 | 1 | 0.03% |
| 10 | 4 | 1 | 0.03% |
| 4 | 10 | 1 | 0.03% |
| 1 | 13 | 1 | 0.03% |
| 8 | 6 | 1 | 0.03% |
| 12 | 1 | 1 | 0.03% |
| 3 | 10 | 1 | 0.03% |
| 7 | 6 | 1 | 0.03% |
| 5 | 8 | 1 | 0.03% |
| 4 | 9 | 1 | 0.03% |
| 11 | 1 | 1 | 0.03% |
| 1 | 11 | 1 | 0.03% |
| 5 | 7 | 1 | 0.03% |
| 2 | 9 | 1 | 0.03% |

## References

1. Yang X, Wang Y, Byrne R, Schneider G, Yang S. Concepts of Artificial Intelligence for Computer-Assisted Drug Discovery. Chem Rev. 2019, 119, 10520–10594

2. Zhu H. Big Data and Artificial Intelligence Modeling for Drug Discovery. Annu Rev Pharmacol Toxicol. 2020, 60, 573–589.

3. Huang K, Fu T, Gao W, Zhao Y, Roohani Y, Leskovec J, Coley CW, Xiao C, Sun J, Zitnik M. Artificial intelligence foundation for therapeutic science. Nat Chem Biol. 2022, 18, 1033–1036

4. Bolcato G, Cuzzolin A, Bissaro M, Moro S, Sturlese M. Can We Still Trust Docking Results? An Extension of the Applicability of DockBench on PDBbind Database. Int J Mol Sci. 2019, 20, 3558.

5. Vittorio S, Lunghini F, Morerio P, Gadioli D, Orlandini S, Silva P, Jan Martinovic, Pedretti A, Bonanni D, Del Bue A, Palermo G, Vistoli G, Beccari AR. Addressing docking pose selection with structure-based deep learning: Recent advances, challenges and opportunities. Comput Struct Biotechnol J. 2024, 23, 2141–2151

6. Cleves AE, Jain AN. Structure- and Ligand-Based Virtual Screening on DUD-E+: Performance Dependence on Approximations to the Binding Pocket. J Chem Inf Model. 2020, 60, 4296–4310.

7. Pedretti A, Mazzolari A, Gervasoni S, Vistoli G. Rescoring and Linearly Combining: A Highly Effective Consensus Strategy for Virtual Screening Campaigns. Int J Mol Sci. 2019, 20, 2060

8. Wang R., Fang X., Lu Y., Wang S. The PDBbind database: collection of binding affinities for protein−ligand complexes with known three-dimensional structures. J Med Chem. 2004, 47, 2977–2980.

9. Wang, R.; Fang, X.; Lu, Y.; Yang, C.-Y.; Wang, S. “The PDBbind Database: Methodologies and updates”, J. Med. Chem., 2005, 48, 4111–4119.

10. https://www.pdbbind-plus.org.cn

11. Zhang C, Zhang X, Freddolino L, Zhang Y. BioLiP2: an updated structure database for biologically relevant ligand-protein interactions. Nucleic Acids Res. 2024, 52(D1), D404–D412

12. Durairaj Janani, Adeshina Yusuf, Cao Zhonglin, Zhang Xuejin, Oleinikovas Vladas, Duignan Thomas, McClure Zachary, Robin Xavier, Kovtun Danny, Rossi Emanuele, Zhou Guoqing, Veccham Srimukh, Isert Clemens, Peng Yuxing, Sundareson Prabindh, Akdel Mehmet, Corso Gabriele, Stärk Hannes, Carpenter Zachary, Bronstein Michael, Kucukbenli Emine, Schwede Torsten, Naef Luca, PLINDER: The protein-ligand interactions dataset and evaluation resource, bioRxiv, 2024.07.17.603955

13. Rohrer SG, Baumann K. Maximum unbiased validation (MUV) data sets for virtual screening based on PubChem bioactivity data. J Chem Inf Model. 2009, 49, 169–84

14. Huang N, Shoichet BK, Irwin JJ. Benchmarking sets for molecular docking. J Med Chem. 2006, 49, 6789–801

15. Mysinger MM, Carchia M, Irwin JJ, Shoichet BK. Directory of useful decoys, enhanced (DUD-E): better ligands and decoys for better benchmarking. J Med Chem. 2012, 55, 6582–94

16. Zhang X, Shen C, Liao B, Jiang D, Wang J, Wu Z, Du H, Wang T, Huo W, Xu L, Cao D, Hsieh CY, Hou T. TocoDecoy: A New Approach to Design Unbiased Datasets for Training and Benchmarking Machine-Learning Scoring Functions. J Med Chem. 2022, 65, 7918–7932

17. Tran-Nguyen VK, Jacquemard C, Rognan D. LIT-PCBA: An Unbiased Data Set for Machine Learning and Virtual Screening. J Chem Inf Model. 2020, 60, 4263–4273

18. Zhou Y, Zhang Y, Zhao D, Yu X, Shen X, Zhou Y, Wang S, Qiu Y, Chen Y, Zhu F. TTD: Therapeutic Target Database describing target druggability information. Nucleic Acids Res. 2024, 52(D1), D1465–D1477

19. https://idrblab.net/ttd/

20. Waterhouse A, Bertoni M, Bienert S, Studer G, Tauriello G, Gumienny R, Heer FT, de Beer TAP, Rempfer C, Bordoli L, Lepore R, Schwede T. SWISS-MODEL: homology modelling of protein structures and complexes. Nucleic Acids Res. 2018, 46(W1), W296–W303

21. Varadi M, Anyango S, Deshpande M, Nair S, Natassia C, Yordanova G, Yuan D, Stroe O, Wood G, Laydon A, Žídek A, Green T, Tunyasuvunakool K, Petersen S, Jumper J, Clancy E, Green R, Vora A, Lutfi M, Figurnov M, Cowie A, Hobbs N, Kohli P, Kleywegt G, Birney E, Hassabis D, Velankar S. AlphaFold Protein Structure Database: massively expanding the structural coverage of protein-sequence space with high-accuracy models. Nucleic Acids Res. 2022, 50(D1), D439–D444.

22. Pedretti A, Mazzolari A, Gervasoni S, Fumagalli L, Vistoli G. The VEGA suite of programs: an versatile platform for cheminformatics and drug design projects. Bioinformatics. 2021, 37, 1174–1175.

23. Meng EC, Goddard TD, Pettersen EF, Couch GS, Pearson ZJ, Morris JH, Ferrin TE. UCSF ChimeraX: Tools for structure building and analysis. Protein Sci. 2023, 32, e4792

24. Webb B, Sali A. Protein Structure Modeling with MODELLER. Methods Mol Biol. 2021, 2199, 239–255

25. Phillips JC, Hardy DJ, Maia JDC, Stone JE, Ribeiro JV, Bernardi RC, Buch R, Fiorin G, Hénin J, Jiang W, McGreevy R, Melo MCR, Radak BK, Skeel RD, Singharoy A, Wang Y, Roux B, Aksimentiev A, Luthey-Schulten Z, Kalé LV, Schulten K, Chipot C, Tajkhorshid E. Scalable molecular dynamics on CPU and GPU architectures with NAMD. J Chem Phys. 2020, 153, 044130

26. Campanella JJ, Bitincka L, Smalley J. MatGAT: an application that generates similarity/identity matrices using protein or DNA sequences. BMC Bioinformatics. 2003, 4, 29

27. Yang J., Roy A., Zhang Y.. Protein-ligand binding site recognition using complementary binding-specific substructure comparison and sequence profile alignment. Bioinformatics. 2013, 29, 2588–2595

28. Zhang Y, Skolnick J. TMalign: a protein structure alignment algorithm based on the TM-score. Nucleic Acids Res. 2005, 33, 2302–9.

29. L. Kaufman and P.J. Rousseeuw, Finding Groups in Data: An Introduction to Cluster Analysis, pp. 68–125, John Wiley and Sons, New Jersey, 2005.

30. E. Schubert, L. Lenssen, Fast k-medoids Clustering in Rust and Python, J. Open Source Softw. 2022, 7, 4183

31. Zdrazil B, Felix E, Hunter F, Manners EJ, Blackshaw J, Corbett S, de Veij M, Ioannidis H, Lopez DM, Mosquera JF, Magarinos MP, Bosc N, Arcila R, Kizilören T, Gaulton A, Bento AP, Adasme MF, Monecke P, Landrum GA, Leach AR. The ChEMBL Database in 2023: a drug discovery platform spanning multiple bioactivity data types and time periods. Nucleic Acids Res. 2024, 52(D1), D1180–D1192

32. Kim S, Chen J, Cheng T, Gindulyte A, He J, He S, Li Q, Shoemaker BA, Thiessen PA, Yu B, Zaslavsky L, Zhang J, Bolton EE. PubChem 2023 update. Nucleic Acids Res. 2023, 51(D1), D1373–D1380

33. https://www.rdkit.org

34. Stewart JJ. Optimization of parameters for semiempirical methods VI: more modifications to the NDDO approximations and re-optimization of parameters. J Mol Model. 2013, 19, 1–32

35. D. Rogers, M. Hahn, Extended-connectivity fingerprints, J Chem Inf Model, 2010, 50, 742–754

36. van der Maaten, L.J.P.; Hinton, G.E. Visualizing Data Using t-SNE *J*. Mach. Learn. Res. 2008, 9, 2579–2605

37. https://scikit-learn.org/stable/

38. G.W. Bemis, M.A. Murcko, The properties of known drugs. 1. Molecular frameworks, J. Med. Chem. 1996, 39, 2887–2893

39. https://icd.who.int/en/

40. Krissinel E. On the relationship between sequence and structure similarities in proteomics. Bioinformatics. 2007, 23, 717–23

41. https://3dmol.org/

